# Capturing early events in aryl hydrocarbon receptor activation using two complementary protein-protein interaction assays

**DOI:** 10.64898/2026.06.19.732932

**Authors:** Tim Kühn, Sarka Tumova, Nicholas Zacharewski, Paul Johannes Averdung, Bianca Berdel, Karl-Heinz Kellner, Stefan Pusch, Marek Jindra, Christiane A. Opitz, Mirja Tamara Prentzell

## Abstract

The aryl hydrocarbon receptor (AHR) is a ligand-activated transcription factor that enables cellular adaptation to environmental, nutritional and metabolic cues. Upon ligand binding, AHR translocates to the nucleus, heterodimerizes with the AHR nuclear translocator (ARNT) and regulates gene expression. Current approaches to measure AHR activity rely on transcriptional readouts, which vary depending on cell type and ligand. Here, we introduce two complementary protein-protein interaction-based assays that detect AHR activation by monitoring AHR-ARNT complex formation. Split-luciferase (NanoBiT) and bimolecular fluorescence complementation (BiFC) detect AHR activation independently of transcriptional output, capturing agonist- and antagonist-dependent AHR modulation across multiple ligands and cellular contexts. NanoBiT enables rapid, real-time analysis of AHR dimerization, whereas BiFC supports imaging of AHR interactions at subcellular resolution. The assays capture further attributes of AHR signaling, including dissociation from chaperones or HIF-1α-mediated competition for ARNT, and enable detection of AHR activation in biological samples. Hence, both assays provide versatile tools to study AHR signaling.

## Background

The aryl hydrocarbon receptor (AHR) is a central integrator of environmental, metabolic and microbial cues that shape cellular behavior across tissues. By sensing chemically diverse endogenous, dietary and xenobiotic ligands, AHR links external exposures and internal metabolic states to transcriptional programs that govern immunity, barrier function, cellular differentiation and stress responses^1,2^. This positions AHR at the interface of environmental health, immunology, metabolism, microbiome research and cancer biology^3,4^. Dysregulated AHR signaling has been implicated in cancer, inflammatory disorders and immune dysfunction, motivating efforts to modulate this pathway for therapeutic purposes. Both AHR agonists and antagonists have progressed into advanced preclinical development and clinical evaluation across multiple indications^5^. At the same time, the pronounced ligand- and context-dependence of AHR signaling underscores the need for experimental approaches that directly and quantitatively assess AHR activation across diverse cellular and disease settings^5^.

AHR belongs to the family of transcription factors characterized by the presence of basic helix-loop-helix (bHLH) and PAS domains, where PAS refers to <u>P</u>ER (period), <u>A</u>RNT (AHR nuclear translocator), and <u>S</u>IM (single-minded)^6–8^. Its modular structure comprises the N-terminal bHLH domain involved in DNA recognition and dimerization, two PAS domains mediating heterodimerization (PAS-A) and ligand binding (PAS-B), and a largely unstructured C-terminal transactivation domain^7,9^. In the absence of ligands, AHR resides in the cytoplasm in a multiprotein complex containing a heat shock protein 90 (HSP90) dimer^10^, AHR-interacting protein (AIP/XAP2)^11^ and prostaglandin E synthase 3 (PTGES3/p23)^12^. Ligand binding induces conformational changes that promote AHR dissociation from cytoplasmic chaperones, AHR nuclear translocation, heterodimerization with the bHLH-PAS partner ARNT, and binding to xenobiotic response elements (XREs), resulting in transcriptional activation of AHR target genes^13^. Coordinated interactions between the bHLH and both PAS domains of AHR and ARNT are required for functional heterodimerization^14,15^.

Quantification of AHR activity remains challenging, as transcriptional outputs can vary substantially depending on ligand identity, cell type, exposure duration and metabolic context^16^. Transcriptomic approaches have provided important insights into global AHR activity signatures and ligand-specific transcriptional programs^17^, but their complexity may limit routine use in experimental or screening settings. Consequently, AHR activity is most commonly assessed using established transcription-based assays, including XRE-driven luciferase reporters and quantitative real-time polymerase chain reaction (qRT-PCR) analysis of canonical AHR target genes such as cytochrome P450 family 1 subfamily A member 1 and subfamily B member 1 (*CYP1A1* and *CYP1B1*)^18,19^. These approaches have been instrumental in advancing the field and remain widely used standards. However, transcriptional readouts represent indirect measures of AHR activation and may not fully capture ligand-specific or context-dependent features of AHR signaling. In addition, ligand-induced proteasomal degradation of AHR uncouples receptor abundance from transcriptional output^20,21^, adding yet another layer of complexity to data interpretation. Together, these considerations motivated the development of protein-centered approaches that directly monitor AHR activation and complement the established transcriptional assays.

In this study, we establish two protein-centered approaches, the NanoLuc Binary Technology (NanoBiT) and bimolecular fluorescence complementation (BiFC), as complementary assays for direct detection and quantification of AHR activity based on its interaction with ARNT. NanoBiT employs reconstitution of a luciferase^22^, while BiFC relies on the reconstitution of a fluorescent protein^23^ upon the interaction of two proteins. Given that heterodimerization of AHR with ARNT represents the defining step in canonical AHR signaling, monitoring this protein-protein interaction offers a direct and mechanistically grounded readout of AHR activation independent of downstream transcriptional responses.

We demonstrate that both assays capture agonist- and antagonist-dependent modulation of AHR signaling across multiple ligands, concentrations and cellular contexts. Benchmarking NanoBiT and BiFC against established qRT-PCR- and XRE-driven reporter assays highlights how protein interaction-based readouts can complement existing methodologies by directly assessing AHR activation. In addition to monitoring AHR-ARNT heterodimerization, the assays resolved ligand-dependent dissociation of AHR from chaperones, thereby interrogating early events within the AHR activation cascade. Moreover, the assays enable investigation of the crosstalk between AHR and other signaling pathways. Furthermore, the detection of AHR activation by stool samples highlights their applicability to biologically complex settings. Together, these findings establish NanoBiT and BiFC as versatile tools for investigation of AHR signaling across a range of mechanistic and physiological contexts.

## Results

### Establishment of NanoBiT and BiFC to measure AHR-ARNT interaction

To directly monitor AHR activation independently of transcriptional output, we developed two complementary protein interaction-based assays that exploit ligand-induced heterodimerization of AHR with ARNT. In the NanoBiT assay, two luciferase fragments were attached to AHR and ARNT, enabling luminescence-based detection of their interaction (**Fig. 1a**). Similarly, the BiFC assay employs AHR and ARNT fused to complementary fragments of a red fluorescent protein (**Fig. 1b**), allowing visualization and quantification of heterodimerization by fluorescence microscopy, for which we developed a custom ImageJ-based analysis workflow (**Extended Data Fig. 1a-c**). To validate both systems, cells were stimulated with the AHR agonists 6-formylindolo[3,2-b]carbazole (FICZ)^24^ and indirubin^25^, two high-affinity indole-derived ligands that differ in their origin and kinetics and are routinely used to activate AHR-dependent gene expression in experimental systems. In cells expressing the NanoBiT fusion constructs (**Extended Data Fig. 2a**), both agonists induced a rapid increase in luminescence relative to DMSO-treated controls, consistent with AHR-ARNT heterodimerization within minutes of ligand exposure (**Fig. 1c**). Quantification across independent experiments demonstrated reproducible enhancement of the luminescence signal (**Fig. 1d**).

**Fig. 1.**
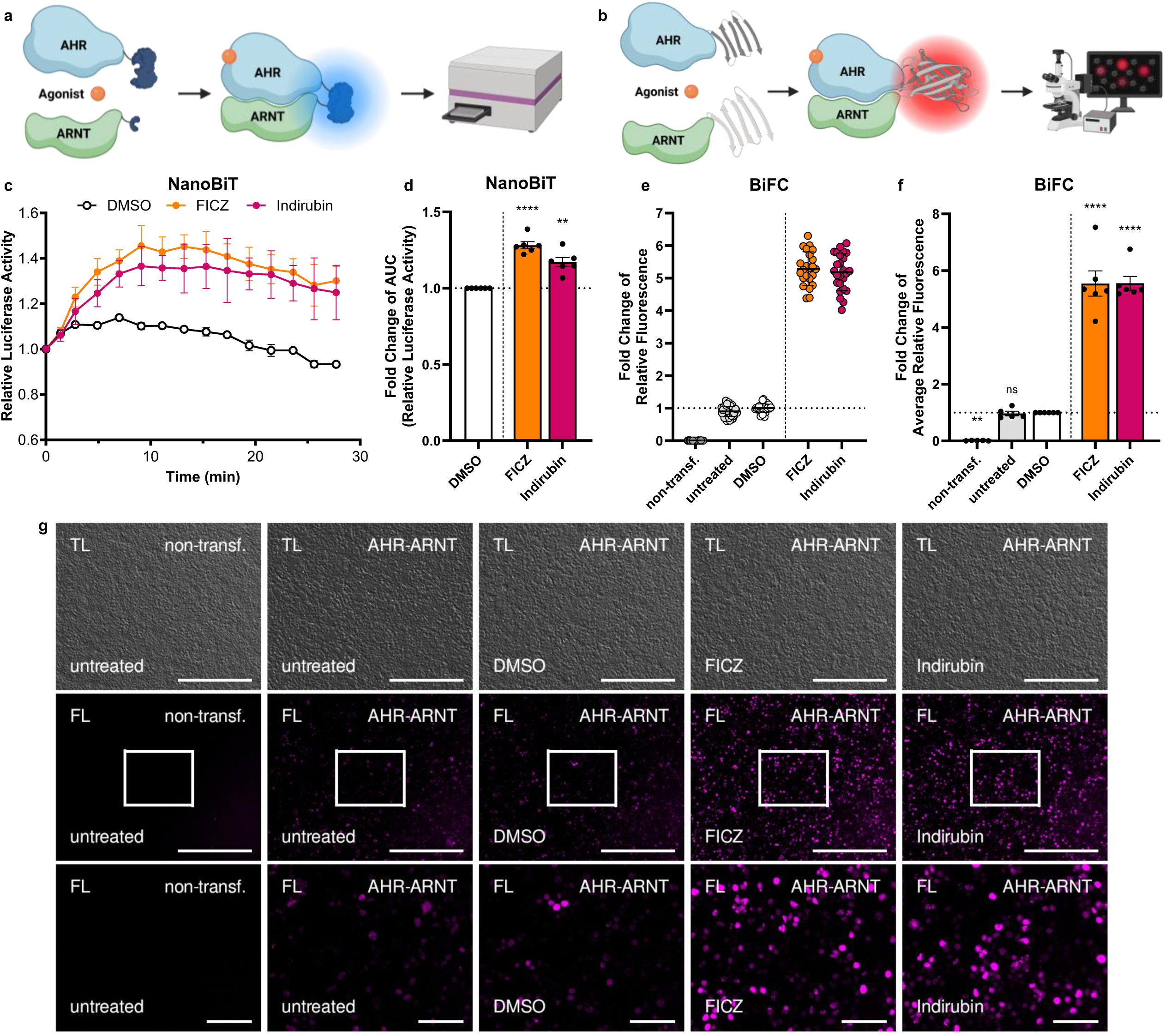
Establishment of NanoBiT and BiFC assays to measure AHR-ARNT interaction. **a,** Schematic of the NanoBiT assay. AHR and ARNT were fused to the large (LgBiT) and small (SmBiT) fragments of NanoLuc luciferase, respectively. Ligand-induced AHR-ARNT interaction reconstitutes luciferase activity, which is detected as luminescence using a plate reader. Image created with BioRender. **b,** Schematic of the bimolecular fluorescence complementation (BiFC) assay. AHR and ARNT were fused to complementary fragments of a red fluorescent protein. Ligand-induced interaction results in fluorescence complementation that is detected by fluorescence microscopy. Image created with BioRender. **c,** Representative NanoBiT time-course in CHO cells showing relative luciferase activity upon treatment with 5 nM FICZ or indirubin compared to DMSO control. Data are shown as mean ± SD of two technical replicates. **d,** Quantification of NanoBiT data (example shown in **c**). The area under the curve (AUC) was calculated for each condition and expressed as fold change relative to DMSO (set to 1). Data are shown as mean ± SEM of n = 6 independent biological replicates. Statistical significance was assessed using one-sample *t*-test on log-transformed fold-change values, followed by Holm-Šidák correction for multiple comparisons. *p* = 6.46 × 10⁻L for FICZ (****); *p* = 1.69 × 10⁻³ for indirubin (**). **e,** Representative BiFC experiment in HEK293T cells showing normalized fluorescence-positive area upon treatment with 100 nM FICZ or indirubin, expressed as fold change relative to DMSO (average = 1). Each dot represents a single image (with an independent, non-overlapping field of view). Non-transfected and untreated cells are shown as controls. Data are shown as mean ± SD of n ≥ 22 images per condition. **f,** Quantification of BiFC fluorescence (example shown in **e**). The average fluorescence per condition was calculated and normalized to DMSO (set to 1). Data are shown as mean ± SEM of n ≥ 5 independent biological replicates. Statistical significance was assessed using one-sample *t*-test on log-transformed fold-change values, followed by Holm-Šidák correction for multiple comparisons. *p* = 1.49 × 10⁻³ to non-transfected (**); *p* = 0.57 to untreated (ns); *p* = 1.04 × 10⁻L for FICZ (****); *p* = 5.86 × 10⁻L for indirubin (****). **g,** Representative BiFC microscopy images corresponding to **e**. Top, transmitted light (TL); middle, fluorescence (FL); bottom, magnified views of the boxed regions in the FL images. Fluorescence signals are pseudocolored in magenta for visualization. Scale bars, 500 µm (top and middle panels) and 100 µm (bottom panel).

To further confirm the specific dependence of the NanoBiT assay on agonist binding, we introduced a point mutation within the ligand-binding PAS-B domain of AHR. Specifically, tyrosine 322 was substituted with alanine (Y322A), targeting a residue previously shown to be required for ligand binding^26^. Indeed, in contrast to wild-type AHR (**Fig. 1c**), the Y322A mutant failed to induce luminescence upon stimulation with FICZ or indirubin, indicating loss of ligand-induced heterodimerization (**Extended Data Fig. 2b-e**). Immunoblot analysis confirmed comparable expression levels of WT and mutant AHR constructs (**Extended Data Fig. 2f**), demonstrating that the observed loss of signal was not attributable to differences in protein abundance. Hence, we conclude that the NanoBiT signal specifically reflects ligand-dependent AHR-ARNT interaction.

Similar to NanoBiT, expression of the BiFC fusion proteins (**Extended Data Fig. 2g**) in combination with agonist treatment induced a robust fluorescent signal, whereas untreated or DMSO-treated cells displayed only minimal fluorescence (**Fig. 1e-g**), consistent with low basal AHR activity from endogenous media-derived ligands^27,28^. Fluorescent complementation was observed exclusively in cells co-transfected with both AHR and ARNT constructs and was absent from cells transfected with control vectors (**Extended Data Fig. 2h, i**), demonstrating that the signal specifically reflected AHR-ARNT interaction rather than non-specific fragment association. Co-staining with DAPI revealed predominant nuclear localization of the BiFC signal, in line with the notion that the AHR-ARNT interaction occurs in the nucleus (**Extended Data Fig. 2j**)^29^. Together, these data establish NanoBiT and BiFC as robust protein-based assays for detecting ligand-induced AHR-ARNT interaction.

### Concentration-dependent AHR activation by FICZ and indirubin

We next evaluated the dynamic range and sensitivity of both assays. Treatment with increasing concentrations of FICZ (**Fig. 2a-e**) or indirubin (**Fig. 2f-j**) resulted in a dose-dependent induction of AHR-ARNT interaction as measured in either assay. From these data, we also calculated the half maximal effective concentration (EC_50_) of both ligands (**Extended Data Fig. 3a-d**). Indirubin was about 6-fold more potent in both assays with EC_50_ of approximately 15 pM (NanoBiT) and 90 pM (BiFC), compared to the EC_50_ for FICZ determined as being around 100 pM (NanoBiT) and 500 pM (BiFC). For comparison, we quantified transcriptional induction of the canonical AHR target genes *CYP1A1* and *CYP1B1* by qRT-PCR in the cells previously analyzed by BiFC. qRT-PCR data confirmed the concentration-dependent effects of agonist treatment for both target genes (**Extended Data Fig. 3e-h**). Of note, EC_50_ for FICZ (2400 and 550 pM) and indirubin (400 and 150 pM) based on transcriptional activation of *CYP1A1* and *CYP1B1* (**Extended Data Fig. 3i-l**) were slightly higher than the EC_50_ obtained for the protein interaction assays. Again, indirubin was more effective compared to FICZ, with *CYP1B1* being more responsive towards both agonists. We conclude that both NanoBiT and BiFC enable sensitive and concentration-dependent detection of AHR activation with EC_50_ in the pM range. Hence, they provide orthogonal, transcription-independent measures of AHR activity and are well suited for quantitative agonist profiling.

**Fig. 2.**
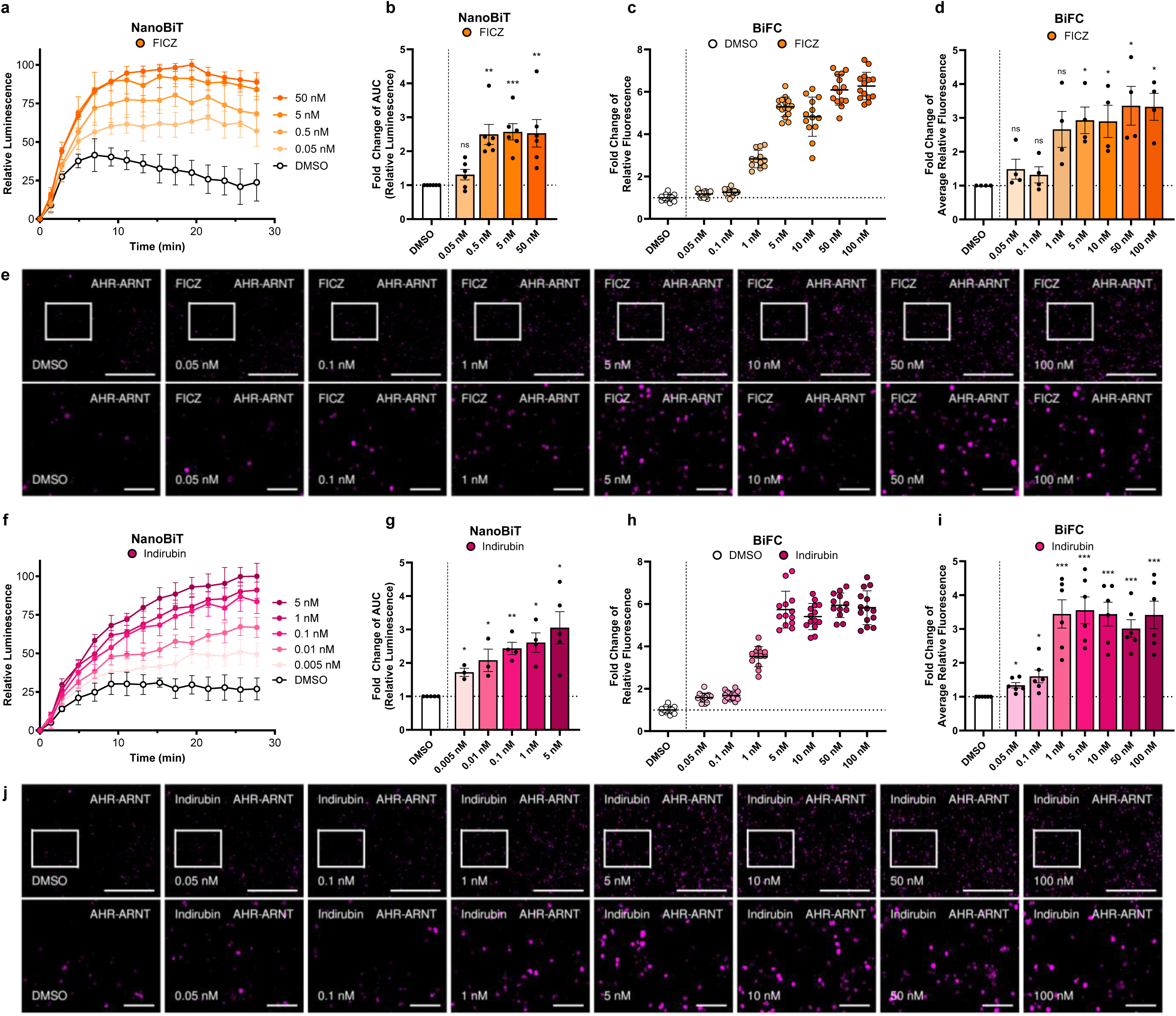
Concentration-dependent detection of AHR activation by NanoBiT and BiFC. **a,** Representative NanoBiT time-course in CHO cells showing relative luminescence following treatment with increasing concentrations of FICZ. Luminescence was normalized to the highest FICZ concentration (50 nM) (set to 100%). Data are shown as mean ± SD of two technical replicates. **b,** Quantification of NanoBiT luminescence data (example shown in **a**). The area under the curve (AUC) was calculated for each condition and expressed as fold change relative to DMSO (set to 1). Data are mean ± SEM of n = 6 independent biological replicates. *p* = 0.12 for 0.05 nM (ns); *p* = 1.00 × 10⁻³ for 0.5 nM (**); *p* = 6.83 × 10 ^4^ for 5 nM (***); *p* = 4.18 × 10 ^3^ for 50 nM (**). **c,** Representative BiFC experiment in HEK293T cells showing normalized fluorescence-positive area upon treatment with increasing concentrations of FICZ, expressed as fold change relative to DMSO (average = 1). Each dot represents a single image (with an independent, non-overlapping field of view). Data are shown as mean ± SD of n ≥ 12 images per condition. **d,** Quantification of BiFC fluorescence (example shown in **c**). The average fluorescence per condition was calculated and normalized to DMSO (set to 1). Data are mean ± SEM of n = 4 independent biological replicates. *p* = 0.27 for 0.05 nM FICZ (ns); *p* = 0.28 for 0.1 nM FICZ (ns); *p* = 5.53 × 10 ^2^ for 1 nM FICZ (ns); *p* = 2.17 × 10 ^2^ for 5 nM FICZ (*); *p* = 3.44 × 10 ^2^ for 10 nM FICZ (*); *p* = 2.99 × 10 ^2^ for 50 nM FICZ (*); *p* = 2.02 × 10 ^2^ for 100 nM FICZ (*). **e,** Representative BiFC fluorescence microscopy images corresponding to **c**. Top, uncropped fluorescence images; bottom, magnified views of the boxed regions shown above. Fluorescence signals are pseudocolored in magenta for visualization. Scale bars, 500 µm (top panel) and 100 µm (bottom panel). **f,** Representative NanoBiT time-course in CHO cells showing relative luminescence following treatment with increasing concentrations of indirubin. Luminescence was normalized to the highest indirubin concentration (5 nM) (set to 100%). Data are shown as mean ± SD of two technical replicates. **g,** Quantification of NanoBiT luminescence data (example shown in **f**). AUC values were calculated and expressed as fold change relative to DMSO (set to 1). Data are mean ± SEM of n ≥ 3 independent biological replicates. *p* = 3.34 × 10 ^2^ for 0.005 nM (*); *p* = 4.46 × 10 ^2^ for 0.01 nM (*); *p* = 6.25 × 10 ^3^ for 0.1 nM (**); *p* = 1.36 × 10 ^2^ for 1 nM (*); *p* = 1.36 × 10 ^2^ for 5 nM (*). **h,** Representative BiFC experiment in HEK293T cells showing normalized fluorescence-positive area upon treatment with increasing concentrations of indirubin, expressed as fold change relative to DMSO (average = 1). Each dot represents a single image (with an independent, non-overlapping field of view). Data are shown as mean ± SD of n ≥ 12 images per condition. **i,** Quantification of BiFC fluorescence (example shown in **h**). The average fluorescence per condition was calculated and normalized to DMSO (set to 1). Data are mean ± SEM of n = 6 independent biological replicates. *p* = 1.16 × 10 ^2^ for 0.05 nM indirubin (*); *p* = 1.16 × 10 ^2^ for 0.1 nM indirubin (*); *p* = 8.05 × 10 ^4^ for 1 nM indirubin (***); *p* = 6.71 × 10 ^4^ for 5 nM indirubin (***); *p* = 6.71 × 10 ^4^ for 10 nM indirubin (***); *p* = 3.40 × 10 ^4^ for 50 nM indirubin (***); *p* = 7.92 × 10 ^4^ for 100 nM indirubin (***). **j,** Representative BiFC fluorescence microscopy images corresponding to **h**. Top, uncropped fluorescence images; bottom, magnified views of the boxed regions shown above. Fluorescence signals are pseudocolored in magenta for visualization. Scale bars, 500 µm (top panel) and 100 µm (bottom panel). Statistical significance was assessed using one-sample *t*-test on log-transformed fold-change values, followed by Holm-Šidák correction for multiple comparisons (**b**, **d**, **g**, **i**).

### Inhibition of AHR-ARNT interaction by the AHR antagonist KYN-101

To determine whether the NanoBiT and BiFC assays are suitable for quantifying AHR antagonism, we examined the effect of KYN-101, a pyrazolo[1,5-a]pyrimidine-derived AHR antagonist^30^. Cells were stimulated with FICZ or indirubin in the presence of increasing concentrations of KYN-101, and AHR-ARNT interaction was monitored. NanoBiT measurements showed a marked reduction in luminescence upon co-treatment with KYN-101 in the nM range (**Fig. 3a,b,f,g**). In the BiFC system, KYN-101 treatment resulted in a progressive loss of AHR-ARNT interaction with increasing antagonist concentrations, approaching baseline levels with low µM concentrations (**Fig. 3c-e,h-j**). Consistent with these observations, KYN-101 reduced agonist-induced transcription of the AHR target genes *CYP1A1* and *CYP1B1* at similar concentrations as in the BiFC assay (**Extended Data Fig. 4a-d**). These findings indicate that direct monitoring of AHR-ARNT heterodimerization provides a reliable measure of antagonist activity, establishing the NanoBiT and BiFC assays as effective tools for quantifying both agonist- and antagonist-mediated modulation of AHR activity.

**Fig. 3.**
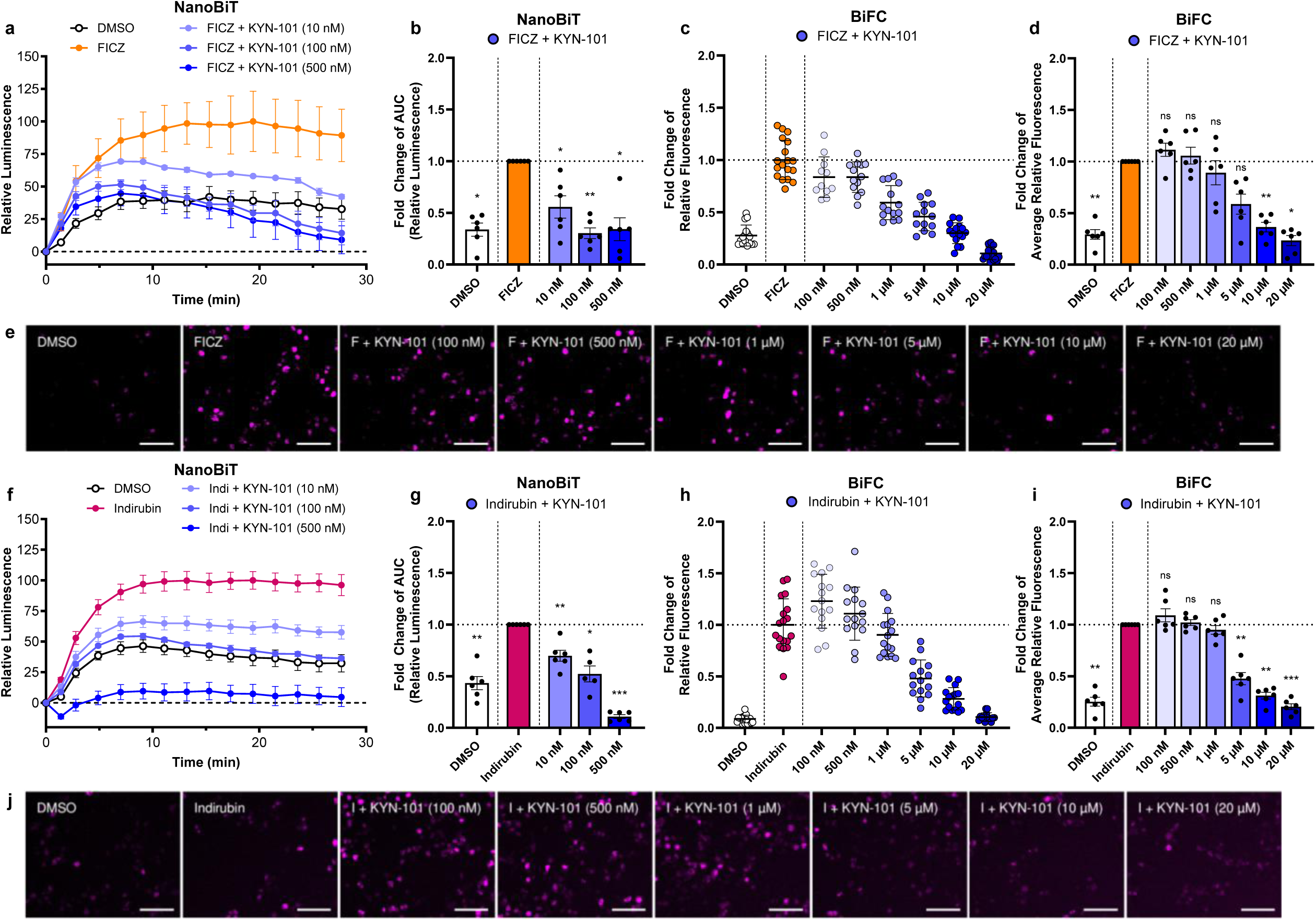
Inhibition of agonist-stimulated AHR-ARNT interaction. **a,** Representative NanoBiT time-course in CHO cells showing relative luminescence following treatment with increasing concentrations of KYN-101 in combination with 5 nM FICZ. Luminescence was normalized to 5 nM FICZ alone (set to 100%). Data are shown as mean ± SD of three technical replicates. **b,** Quantification of NanoBiT luminescence data (example shown in **a**). The area under the curve (AUC) was calculated for each condition and expressed as fold change relative to FICZ alone (set to 1). Data are mean ± SEM of n = 6 independent biological replicates. KYN-101 treatment was performed in presence of 5 nM FICZ. *p* = 3.03 × 10 ^2^ for DMSO (*); *p* = 3.03 × ⁻ for 10 nM KYN-101 (*); *p* = 3.21 × 10⁻ for 100 nM KYN-101 (**); *p* = 3.03 × 10⁻ for 500 nM KYN-101 (*). **c,** Representative BiFC experiment in HEK293T cells showing normalized fluorescence-positive area upon treatment with rising concentrations of KYN-101 in combination with 100 nM FICZ, expressed as fold change relative to FICZ (average = 1). Each dot represents a single image (with an independent, non-overlapping field of view). Data are shown as mean ± SD of n ≥ 11 images per condition. **d,** Quantification of BiFC fluorescence (example shown in **c**). The average fluorescence per condition was calculated and normalized to FICZ alone (set to 1). Data are mean ± SEM of n = 6 independent biological replicates. KYN-101 treatment was performed in presence of 100 nM FICZ. *p* = 6.26 × 10 ^3^ for DMSO (**); *p* = 0.43 for 100 nM KYN-101 (ns); *p* = 0.61 for 500 nM KYN-101 (ns); *p* = 0.52 for 1 µM KYN-101 (ns); *p* = 0.12 for 5 µM KYN-101 (ns); *p* = 4.90 × 10 ^3^ for 10 µM KYN-101 (**); *p* = 1.33 × 10 ^2^ for 20 µM KYN-101 (*). **e,** Representative BiFC fluorescence microscopy images corresponding to **c**. Magnified fluorescence images are shown. Fluorescence signals are pseudocolored in magenta for visualization. Scale bars, 100 µm. **f,** Representative NanoBiT time-course in CHO cells showing relative luminescence in % following treatment with increasing concentrations of KYN-101 in combination with 5 nM indirubin. Luminescence was normalized to 5 nM indirubin alone (set to 100). Data are shown as mean ± SD of three technical replicates. **g,** Quantification of NanoBiT luminescence data (example shown in **f**). AUC values were calculated and expressed as fold change relative to indirubin alone (set to 1). Data are mean ± SEM of n ≥ 5 independent biological replicates. KYN-101 treatment was performed in presence of 5 nM indirubin. *p* = 5.54 × 10 ^3^ for DMSO (**); *p* = 8.69 × 10 ^3^ for 10 nM KYN-101 (**); *p* = 1.15 × 10 ^2^ for 100 nM KYN-101 (*); *p* = 2.21 × 10 ^4^ for 500 nM KYN-101 (***). **h,** Representative BiFC experiment in HEK293T cells showing normalized fluorescence-positive area upon treatment with rising concentrations of KYN-101 in combination with 100 nM indirubin, expressed as fold change relative to indirubin (average = 1). Each dot represents a single image (with an independent, non-overlapping field of view). Data are shown as mean ± SD of n ≥ 15 images per condition. **i,** Quantification of BiFC fluorescence (example shown in **h**). The average fluorescence per condition was calculated and normalized to indirubin alone (set to 1). Data are mean ± SEM of n = 6 independent biological replicates. KYN-101 treatment was performed in presence of 100 nM indirubin (n = 4) or 10 nM indirubin (n = 2). *p* = 4.96 × 10 ^3^ for DMSO (**); *p* = 0.52 for 100 nM KYN-101 (ns); *p* = 0.54 for 500 nM KYN-101 (ns); *p* = 0.52 for 1 µM KYN-101 (ns); *p* = 7.62 × 10 ^3^ for 5 µM KYN-101 (**); *p* = 1.25 × 10 ^3^ for 10 µM KYN-101 (**); *p* = 8.32 × 10 ^4^ for 20 µM KYN-101 (***). **j,** Representative BiFC fluorescence microscopy images corresponding to **h**. Magnified fluorescence images are shown. Fluorescence signals are pseudocolored in magenta for visualization. Scale bars, 100 µm. Statistical significance was assessed using one-sample *t*-test on log-transformed fold-change values, followed by Holm-Šidák correction for multiple comparisons (**b**, **d**, **g**, **i**).

### Additional AHR agonists induce AHR-ARNT heterodimerization

We next examined two additional tryptophan-derived agonists: kynurenic acid (KynA) and indole-3-pyruvic acid (I3P). KynA is generated through the kynurenine pathway, in which tryptophan is initially converted to kynurenine, followed by transamination of kynurenine to KynA catalyzed by kynurenine aminotransferases (KATs)^31^. I3P is produced by transamination of tryptophan via aminotransferases in host and microbial cells and is also generated through oxidative deamination by interleukin-4-induced-1 (IL4I1), an L-amino-acid oxidase^32^. In addition, IL4I1-dependent I3P production can also indirectly lead to KynA formation ^17^. BiFC analysis revealed significant induction of AHR-ARNT interaction upon treatment with all four agonists – FICZ, indirubin, KynA and I3P (**Extended Data Fig. 5a-c**). Among these ligands, KynA consistently elicited the weakest interaction signal, whereas FICZ, indirubin and I3P induced more pronounced AHR-ARNT complementation. To benchmark BiFC against transcript-based AHR activity measurements, we performed qRT-PCR analysis of *CYP1A1* and *CYP1B1* in identically treated, non-transfected HEK293T cells. *CYP1A1* and *CYP1B1* exhibited similar induction across the four agonists, but the extent of induction was lower compared to the response observed in the BiFC assay (**Extended Data Fig. 5d, e**). We additionally assessed expression of TCDD-inducible poly (ADP-ribose) polymerase (*TIPARP*), another frequently used AHR target gene^33,34^. *TIPARP* transcript levels showed rather weak and variable responses upon agonist treatment (**Extended Data Fig. 5f**). Together, these results highlight the ligand-and target gene-specific variability inherent to transcriptional readouts, underscoring the value of complementary protein interaction-based approaches for monitoring AHR activation across diverse ligands.

We next examined AHR activation in glioblastoma cells as a model for tumor-intrinsic AHR activity^35^. We first assessed transcriptional responses upon ligand treatment by qRT-PCR (**Fig. 4a-c).** While all four ligands significantly enhanced *CYP1A1* transcript levels (**Fig. 4a**), *CYP1B1* was only induced in response to KynA and *TIPARP* expression was not significantly altered by any of the tested AHR ligands (**Fig. 4b,c**), reflecting ligand- and target gene-specific variability. Because XRE-driven luciferase assays are also frequently used to assess AHR signaling, we next evaluated XRE reporter activity under the same experimental conditions. Only minor increases in XRE luciferase were detected following treatment with FICZ, indirubin and I3P, whereas KynA did not measurably alter reporter activity compared to control (**Fig. 4d**). Based on these data alone, AHR activation by tryptophan-derived metabolites would be difficult to infer in glioblastoma cells. Notably, BiFC revealed robust AHR-ARNT interaction in response to all four agonists with a consistently high measurement window (**Fig. 4e-g**), highlighting the capacity of the protein-protein interaction assay to verify AHR activation in contexts with transcriptionally diverse outcomes.

**Fig. 4.**
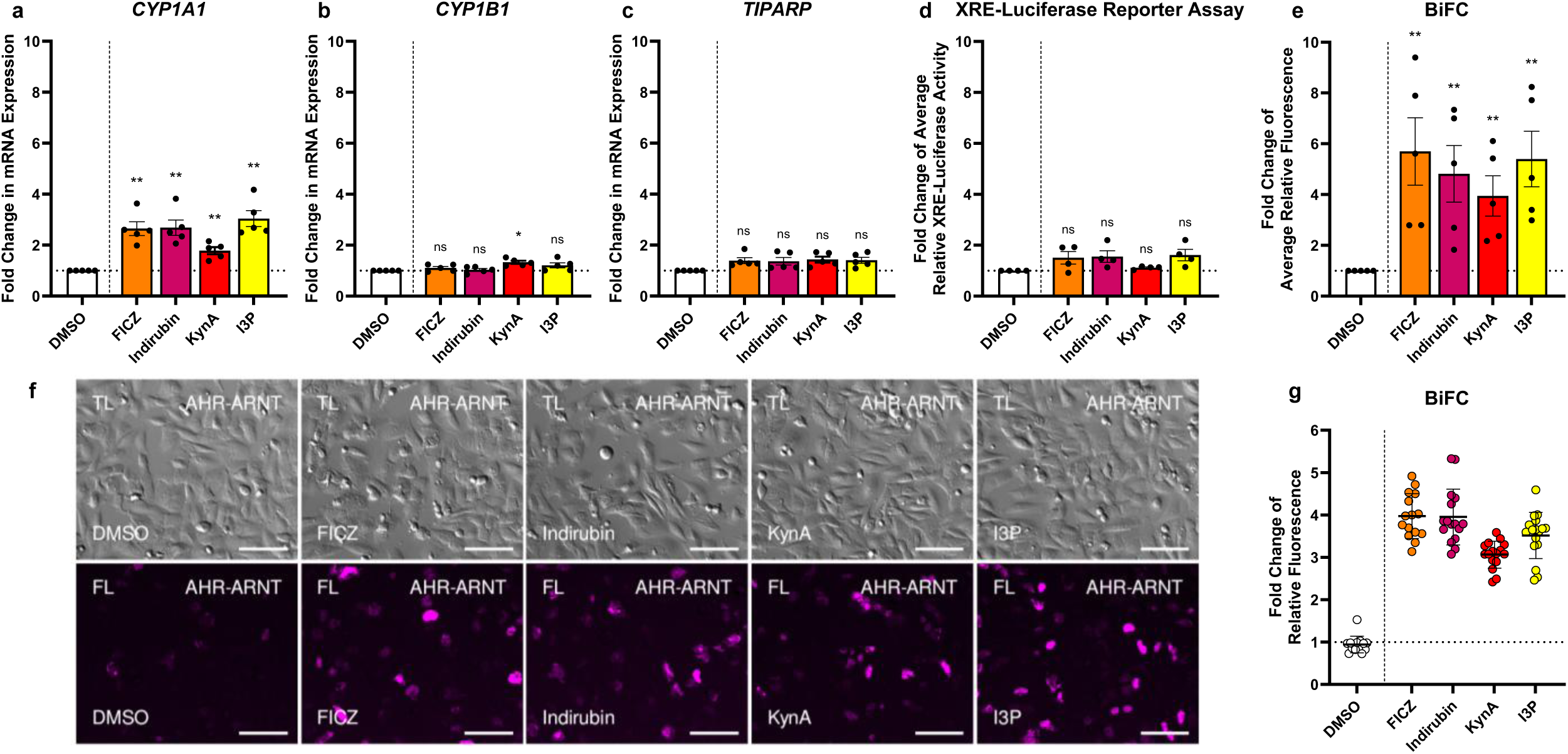
BiFC detects AHR-ARNT interaction in LN-229 cells. **a, b, c,** Quantification of *CYP1A1* (**a**), *CYP1B1* (**b**) and *TIPARP* (**c**) mRNA expression in non-transfected LN-229 cells following agonist treatment, measured by qRT-PCR and expressed as fold change relative to DMSO (set to 1). Data are mean ± SEM of n = 5 independent biological replicates. **a**, *p* = 1.80 × 10 ^3^ for FICZ (**); *p* = 1.80 × 10 ^3^ for indirubin (**); *p* = 2.13 × 10 ^3^ for KynA (**); *p* = 1.40 × 10 ^3^ for I3P (**). **b**, *p* = 0.28 for FICZ (ns); *p* = 0.67 for indirubin (ns); *p* = 1.46 × 10 ^2^ for KynA (*); *p* = 0.25 for I3P (ns). **c**, *p* = 5.17 × 10 ^2^ for FICZ (ns); *p* = 5.17 × 10 ^2^ for indirubin (ns); *p* = 5.17 × 10 ^2^ for KynA (ns); *p* = 5.17 × 10 ^2^ for I3P (ns). **d,** Quantification of XRE-luciferase activity, expressed as fold change relative to DMSO (average luminescence set to 1). Data are mean ± SEM of n = 4 independent biological replicates. *p* = 0.13 for FICZ (ns); *p* = 0.13 for indirubin (ns); *p* = 0.13 for KynA (ns); *p* = 0.13 for I3P (ns). **e,** Quantification of relative BiFC fluorescence. The average fluorescence per condition was calculated and normalized to DMSO (set to 1). Data are mean ± SEM of n = 5 independent biological replicates. *p* = 9.39 × 10 ^3^ for FICZ (**); *p* = 9.39 × 10 ^3^ for indirubin (**); *p* = 9.39 × ⁻ for KynA (**); *p* = 6.01 × 10⁻ for I3P (**). **f,** Representative BiFC fluorescence microscopy images in LN-229 cells corresponding to **g**. Magnified images are shown. Top, transmitted light (TL); bottom, fluorescence (FL). Fluorescence signals are pseudocolored in magenta for visualization. Scale bars, 100 µm. **g,** Representative BiFC experiment in LN-229 cells showing normalized fluorescence-positive area upon treatment with FICZ, indirubin, KynA, or I3P, expressed as fold change relative to DMSO (average = 1). Each dot represents a single image (with an independent, non-overlapping field of view). Data are shown as mean ± SD of n ≥ 14 images per condition. Agonist treatment was conducted with 100 nM FICZ or indirubin, 100 µM KynA or 25 µM I3P. Statistical significance was assessed using one-sample *t*-test on log-transformed fold-change values, followed by Holm-Šidák correction for multiple comparisons (**a-e**).

### Competitive sequestration of ARNT under hypoxia-mimetic conditions

AHR signaling has been proposed to intersect with other transcriptional pathways through competition for shared cofactors, most notably ARNT, which is also known as hypoxia-inducible factor-1β (HIF-1β)^36^. Stabilization of HIF-1α under hypoxic conditions is expected to promote formation of HIF-1α-ARNT complexes and thereby limit the availability of ARNT for AHR heterodimerization (**Fig. 5a**). However, the extent to which such competition occurs, and the cellular contexts, in which it meaningfully impacts AHR signaling, remain incompletely understood^37^. To examine whether pathway-level competition can be captured by protein-interaction assays, we treated cells with dimethyloxalylglycine (DMOG), a hypoxia-mimetic compound that stabilizes HIF-1α by inhibiting prolyl hydroxylases^38^. DMOG treatment markedly reduced the AHR-ARNT interaction as measured by BiFC (**Fig. 5b-d**), consistent with diminished AHR-ARNT complex formation under hypoxia-mimetic conditions resulting from competitive sequestration of ARNT by HIF-1α.

**Fig. 5.**
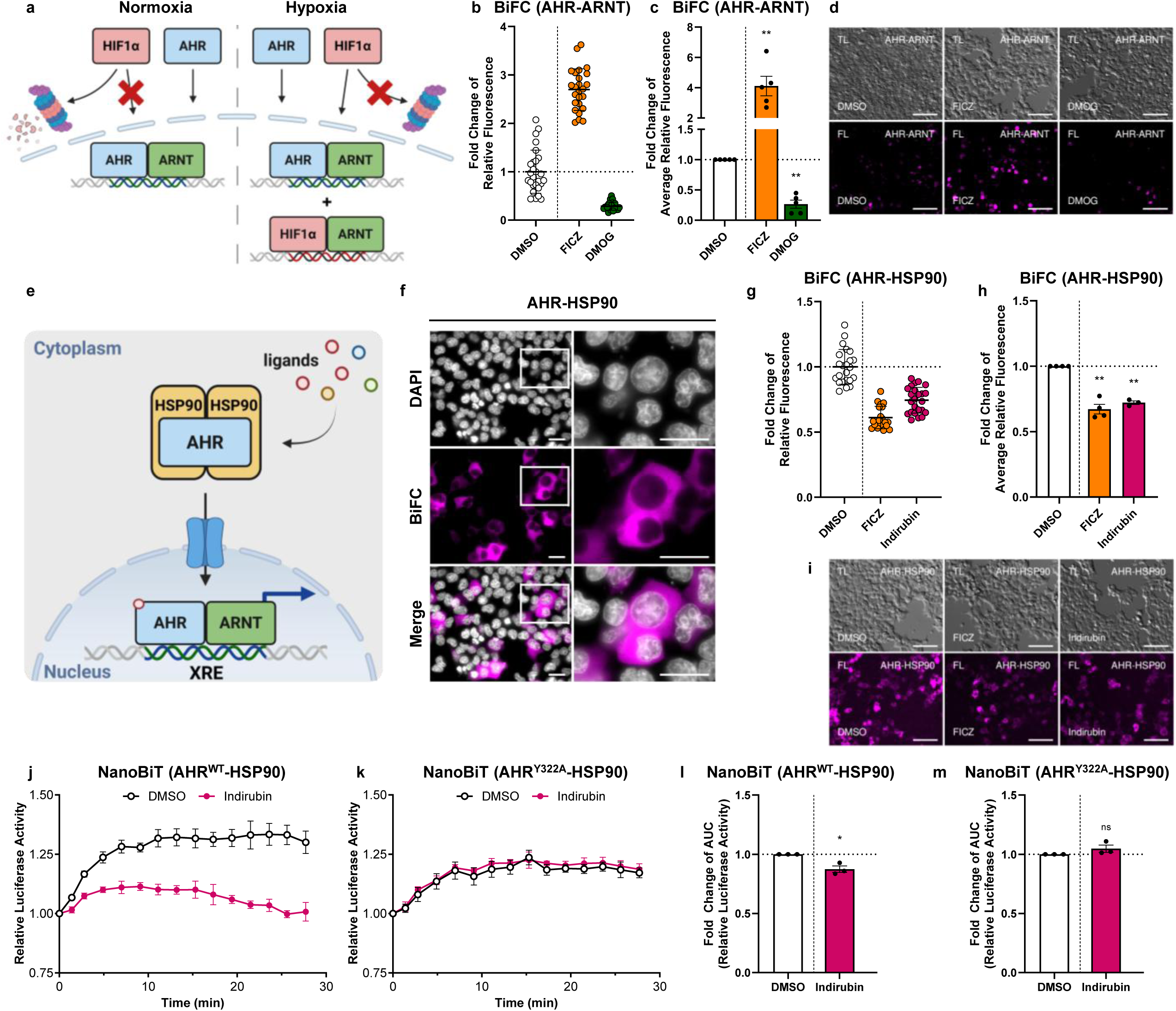
BiFC and NanoBiT enable indirect and direct monitoring of AHR interactions. **a,** Schematic overview of ARNT protein-protein interaction with AHR or HIF1α upon normoxia or hypoxia. Hypoxia stabilizes HIF1α, which competes with AHR for ARNT binding. Image created with BioRender. **b,** Representative BiFC experiment of AHR-ARNT protein-protein interaction in HEK293T cells showing normalized fluorescence-positive area upon treatment with 100 nM FICZ or 2 mM DMOG, expressed as fold change relative to DMSO (average = 1). Each dot represents a single image (with an independent, non-overlapping field of view). Data are shown as mean ± SD of n ≥ 24 images per condition. **c,** Quantification of BiFC fluorescence (example shown in **b**). The average fluorescence per condition was calculated and normalized to DMSO (set to 1). Data are mean ± SEM of n = 5 independent biological replicates. Statistical significance was assessed using one-sample *t*-test on log-transformed fold-change values, followed by Holm-Šidák correction for multiple comparisons. *p* = 1.55 × 10 ^3^ for FICZ (**); *p* = 6.91 × 10 ^3^ for DMOG (**). **d,** Representative BiFC fluorescence microscopy images corresponding to **b**. Magnified images are shown. Top, transmitted light (TL); bottom, fluorescence (FL). Fluorescence signals are pseudocolored in magenta for visualization. Scale bars, 100 µm. **e,** Schematic of AHR interactions in the cytoplasm and in the nucleus. Ligand-binding leads to AHR translocation to the nucleus and dissociation from HSP90. Image created with BioRender. **f,** Representative BiFC microscopy images of AHR-HSP90 protein-protein interaction, combined with nuclear DAPI staining in HEK293T cells. Top, DAPI channel; middle, BiFC channel; bottom, merged images. Enlarged views of the boxed regions in the left panels are shown on the right. DAPI signals are shown in white and BiFC signals are pseudocolored in magenta for visualization. Scale bars, 25 µm. Representative of n = 2 independent biological experiments. **g,** Representative BiFC experiment of AHR-HSP90 protein-protein interaction in HEK293T cells showing normalized fluorescence-positive area upon treatment with 100 nM FICZ or indirubin, expressed as fold change relative to DMSO (average = 1). Each dot represents a single image (with an independent, non-overlapping field of view). Data are shown as mean ± SD of n ≥ 20 images per condition. **h,** Quantification of BiFC fluorescence (example shown in **g**). The average fluorescence per condition was calculated and normalized to DMSO (set to 1). Data are mean ± SEM of n = 4 (FICZ) or n = 3 (indirubin) independent biological replicates. Statistical significance was assessed using one-sample *t*-test on log-transformed fold-change values, followed by Holm-Šidák correction for multiple comparisons. *p* = 6.63 × 10 ^3^ for FICZ (**); *p* = 6.63 × 10 ^3^ for indirubin (**). **i,** Representative BiFC fluorescence microscopy images corresponding to **g**. Magnified images are shown. Top, transmitted light (TL); bottom, fluorescence (FL). Fluorescence signals are pseudocolored in magenta for visualization. Scale bars, 100 µm. **j, k,** Representative NanoBiT time-course of AHR-HSP90 protein-protein interaction using wildtype (AHR^WT^) (**j**) or mutated (AHR^Y322A^) (**k**) AHR construct in CHO cells showing relative luciferase activity upon treatment with 5 nM indirubin compared to DMSO control. Data are shown as mean ± SD of three technical replicates. **l, m,** Quantification of NanoBiT luminescence of wildtype (**j**) and mutated (**k**) AHR. The area under the curve (AUC) was calculated for each condition and expressed as fold change relative to DMSO (set to 1). Data are mean ± SEM of n = 3 independent biological replicates. Statistical significance was assessed using one-sample *t*-test on log-transformed fold-change values. **l**, *p* = 4.96 × 10 ^2^ (*). **m**, *p* = 0.25 (ns).

### Monitoring ligand-induced dissociation of AHR from cytoplasmic chaperones

Beyond direct measurement of AHR-ARNT heterodimerization, both BiFC and NanoBiT enable interrogation of additional protein interactions along the AHR activation cascade. In its inactive state, cytoplasmic AHR is stabilized by association with HSP90 chaperones and co-chaperones, which maintain AHR in a ligand-responsive conformation and protect it from proteasomal degradation^10,39^. Ligand binding induces conformational changes that promote dissociation from HSP90, nuclear translocation and ARNT heterodimerization, although the underlying mechanism has not been fully elucidated (**Fig. 5e**)^40^. We therefore hypothesized that ligand-induced AHR activation would result in reduced AHR-HSP90 interaction measurable by the BiFC or NanoBiT assays with respectively tagged AHR and HSP90 proteins. Consistent with this model, BiFC detected AHR-HSP90 interaction outside the nucleus (**Fig. 5f**), and treatment with FICZ and indirubin significantly decreased the AHR-HSP90 protein-protein interaction (**Fig. 5g-i**). Indirubin led to rapid dissociation of WT AHR from HSP90, as evidenced by a reduction in luminescence signal measured by the NanoBiT assay (**Fig. 5j,l**). In contrast, the AHR Y322A mutant remained associated with HSP90 following ligand exposure (**Fig. 5k,m**), consistent with the notion that the conformational changes required for chaperone release require ligand binding to AHR. Thus, the protein-based assays can be expected to resolve sequential steps of AHR activation, represented here by ligand-dependent dissociation from chaperones.

### Detection of AHR activity as a biomarker of intestinal health

The AHR plays a pivotal role in diverse disease contexts, including cancer, autoimmunity, and inflammatory bowel disease^41^. Within the gastrointestinal tract, the AHR broadly expressed in immune and epithelial cells integrates signals from endogenous, dietary, and microbiota-derived ligands^42^. Gut microbiota and host metabolism produce a broad ensemble of AHR agonists, especially tryptophan catabolites such as indoles and kynurenine derivatives^43^. Impaired microbial conversion of tryptophan into AHR ligands exacerbates mucosal inflammation, whereas ligand-mediated AHR activation promotes epithelial repair and ameliorates inflammation in experimental colitis models^44^. Importantly, net AHR activity in the gut reflects the combined action of multiple metabolites with varied affinities and effects. To assess whether protein-based AHR interaction assays can be applied to measure AHR activity in complex biological matrices such as fecal samples, we treated cells with diluted fecal suspensions obtained from three healthy volunteers (**Fig. 6a-i**). In all cases, the treatment resulted in a marked increase of BiFC-mediated fluorescence, indicating the presence of AHR agonists that enhance AHR-ARNT interaction (**Fig. 6a-i**). These results demonstrate that protein-centered assays detect AHR activity in complex, heterogeneous biological samples where the composition and identity of AHR modulators are unknown.

**Fig. 6.**
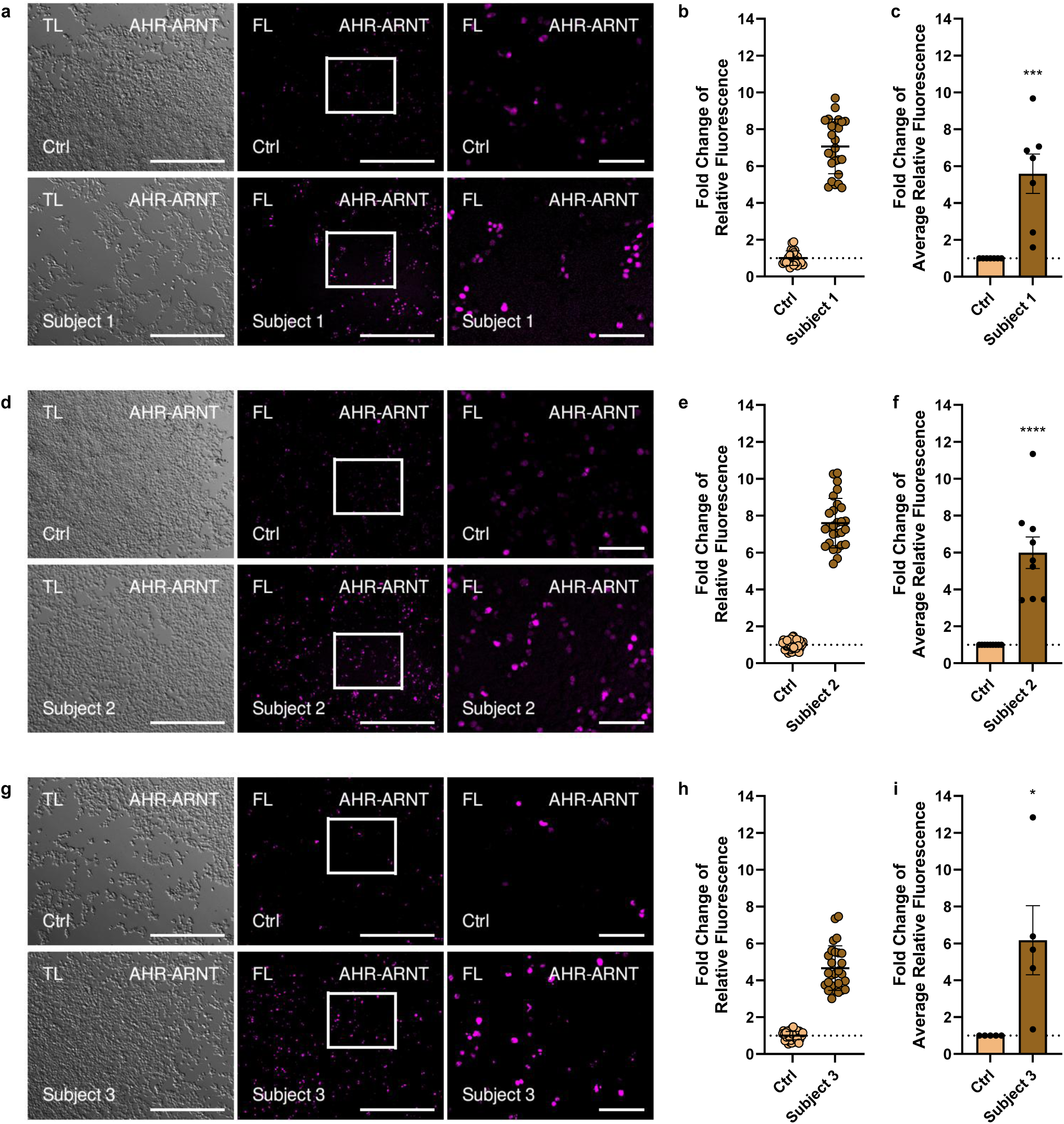
Detection of AHR-activating properties of fecal samples. **a,** Representative BiFC microscopy images in HEK293T cells treated with a diluted stool sample from a healthy donor (subject 1) or PBS as control (Ctrl). Left, transmitted light (TL); middle, fluorescence (FL); right, magnified views of the boxed regions in the FL images. Fluorescence signals are pseudocolored in magenta for visualization. Scale bars, 500 µm (left and middle panels) and 100 µm (right panel). **b,** Representative BiFC experiment in HEK293T cells showing normalized fluorescence-positive area upon treatment with a diluted fecal sample from subject 1, expressed as fold change relative to DMSO (average = 1). Each dot represents a single image (with an independent, non-overlapping field of view). Data are shown as mean ± SD of n ≥ 25 images per condition. **c,** Quantification of BiFC fluorescence corresponding to **b**. The average fluorescence per condition was calculated and normalized to Ctrl (set to 1). Data are mean ± SEM of n = 7 independent biological replicates. Statistical significance was assessed using one-sample *t*-test on log-transformed fold-change values. *p* = 7.06 × 10 ^4^ for subject 1 (***). **d**, Representative BiFC microscopy images of cells treated with a diluted stool sample from a healthy donor (subject 2) as in **a**. **e**, Representative BiFC experiment of cells upon treatment with a diluted fecal sample from subject 2 as in **b**. Data are shown as mean ± SD of n ≥ 18 images per condition. **f**, Quantification of BiFC fluorescence corresponding to **e** and as described in **c**. Data are mean ± SEM of n = 9 independent biological replicates. *p* = 1.76 × 10 ^6^ for subject 2 (****). **g**, Representative BiFC microscopy images of cells treated with a diluted stool sample from a healthy donor (subject 3) as in **a**. **h**, Representative BiFC experiment of cells upon treatment with a diluted fecal sample from subject 3 as in **b**. Data are shown as mean ± SD of n ≥ 27 images per condition. **i**, Quantification of BiFC fluorescence corresponding to **h** and as described in **c**. Data are mean ± SEM of n = 5 independent biological replicates. *p* = 1.23 × 10 ^2^ for subject 3 (*).

## Discussion

Assessment of AHR activity has traditionally relied on target gene expression providing widely used and informative readouts. However, transcriptional readouts depend on a layer of intermediary steps and can substantially vary between cell type, ligand, and experimental context. Here, we introduce NanoBiT and BiFC as protein-based approaches for direct monitoring of AHR-ARNT heterodimerization, the central event in canonical AHR signaling, which offer a direct measure of AHR activation independent of downstream transcriptional variability.

NanoBiT offers superior temporal resolution, enabling real-time monitoring of AHR interaction dynamics within minutes of ligand exposure. BiFC is particularly suited for studying long-term effects of AHR modulation, and enables microscopic visualization of AHR-ARNT interaction, allowing concurrent assessment of cell morphology, viability and subcellular localization of the interaction (**Extended Data Fig. 2j**).

While NanoBiT detected AHR-ARNT interactions with EC_50_ as low as approximately 100 pM for FICZ (**Extended Data Fig. 3a**) and 15 pM for indirubin (**Extended Data Fig. 3b**), BiFC detection of AHR-ARNT interaction required higher concentrations for either agonist, with EC_50_ around 500 pM (FICZ) and 90 pM (indirubin) (**Extended Data Fig. 3c, d**). The shift in EC_50_ and apparent ligand activation thresholds likely reflect the distinct temporal sampling windows of both assays. NanoBiT captures early AHR-ARNT interaction dynamics within minutes of ligand addition (≈30 min), providing a near-instantaneous readout of AHR engagement. In contrast, BiFC measurements were obtained after extended ligand exposure (20-24 h), during which cellular metabolism and CYP-enzyme-mediated feedback regulation^45^ may have altered the effective concentration of AHR ligands, potentially reducing their bioavailability over time. Consequently, BiFC integrates both AHR activation and ligand metabolism, whereas NanoBiT predominantly reflects immediate ligand-induced heterodimerization.

Published studies using yeast AHR assays^25,46^, luciferase reporters^47^, ethoxyresorufin-O-deethylase (EROD) measurements^27,48,49^ as well as competitive binding approaches^50^ demonstrate substantial variability in the reported potencies of classical AHR agonists such as indirubin and FICZ, depending on assay type, species background and exposure duration. Notably, the high (subnanomolar) sensitivity observed in the NanoBiT and BiFC assays (**Extended Data Fig. 3a-d**) is well within the range of established AHR activity assays reported in the literature and supports their suitability as sensitive protein-centered platforms for characterization and screening of AHR ligands, including compounds with comparatively low potency.

Both methods rely on ectopic expression of tagged proteins, which may alter endogenous protein abundance, stoichiometry or other regulatory parameters, potentially affecting subcellular localization and interaction behavior. For NanoBiT, this is mitigated by a design aiming to operate at low expression levels so as to minimize artifacts arising from overexpression or saturated complex formation^22^. For BiFC, the distribution of the exogenously expressed proteins was confirmed by imaging, showing that AHR-HSP90 complexes localized to the cytoplasm (**Fig. 5f**), whereas AHR-ARNT complexes were primarily detected in the nucleus (**Extended Data Fig. 2j**), consistent with the established cytoplasmic and nuclear compartmentalization of their endogenous counterparts. With BiFC providing spatial information on interaction-competent complexes, and NanoBiT monitoring interaction dynamics, the two assays interrogate complementary aspects of AHR protein interactions and their concordant findings support the physiological relevance of our observations.

We demonstrated robust AHR-ARNT interaction induced by diverse tryptophan-derived metabolites, including FICZ, indirubin, KynA, and I3P, across different cell types. These findings underscore the ligand sensitivity of the assays and their applicability in distinct biological contexts. Importantly, direct measurement of AHR-ARNT interaction revealed consistent activation patterns even in settings where classical transcriptional readouts were only modestly induced or were target gene-specific (**Fig 4a-d; Extended Data Fig. 5d-f**). This highlights a key advantage of protein-centered assays in reducing ambiguity due to variability in downstream transcriptional regulation. Such design may be especially relevant when analyzing heterogeneous mixtures, where the identity and relative contribution of individual agonists are typically unknown. Indeed, here we have measured AHR activation by fecal samples (**Fig. 6a-i**), which contain complex and largely undefined mixtures of endogenous and microbial-derived AHR modulators. Because distinct AHR ligands can differentially shape downstream transcriptional programs, gene expression-based readouts may reflect ligand-specific signaling biases rather than overall AHR activation. Protein interaction-based detection instead captures an early and proximal step in AHR activation, enabling assessment of ligand-dependent AHR engagement independently of downstream transcriptional context. This feature may be particularly advantageous for studies involving biologically complex samples, including those derived from clinical or environmental sources, where the heterogeneous ligand composition can complicate interpretation of transcription-based readouts.

While the experiments presented here focus on tryptophan-derived metabolites, the underlying assay principles are not restricted to this ligand class. Given the historical role of AHR as a sensor of environmental xenobiotics^2^, these assays may offer complementary tools for exploratory or screening-based studies aimed at identifying additional classes of AHR ligands, including environmental contaminants.

Beyond agonist detection, both methods also detected pharmacological inhibition of AHR-ARNT interaction by the AHR antagonist KYN-101 (**Fig. 3a-j**). NanoBit measured KYN-101-mediated inhibition in the nM range (**Fig. 3a,b,f,g**), whereas BiFC required low µM concentrations of KYN-101 to achieve significant suppression of AHR-ARNT heterodimerization (**Fig. 3c-e,h-j**). These differences likely reflect the distinct temporal sampling windows of the two assays, with BiFC integrating long-term effects of ligand exposure, consistent with the higher agonist concentrations required in this system. Although KYN-101 has been described by Campesato and colleagues^30^ as having low nanomolar activity in luciferase reporter assays, relatively limited quantitative data are available across alternative assays, emphasizing the value of orthogonal protein interaction-based readouts for monitoring AHR inhibition.

Recent high-resolution cryo-electron microscopy structures of the cytosolic AHR complex have provided important insights into agonist binding^39,40^. However, the detailed molecular mechanisms underlying the structural transition of AHR from a chaperone-bound cytosolic complex to a transcriptionally active nuclear heterodimer remain incompletely understood. A recent computational docking study suggests that AHR antagonists may engage multiple binding sites beyond the orthosteric ligand-binding pocket, including interfaces involving AIP and HSP90^51^, highlighting novel molecular mechanisms of AHR inhibition. In this context, the combined application of NanoBiT or BiFC with site-directed mutagenesis may provide a framework for dissecting ligand-specific modes of AHR modulation, including both agonist- and antagonist-induced interaction dynamics, thereby supporting the mechanistic characterization of AHR activation. Furthermore, both assays can be applied to elucidate other relevant complexes at distinct stages of the AHR signaling pathway, as we show here for the ligand-induced and spatially defined dissociation of the AHR-chaperone complex (**Fig. 5f-m**).

NanoBiT and BiFC enable the investigation of crosstalk between AHR and other transcriptional pathways. Stabilization of HIF-1α under hypoxia-mimetic conditions reduced AHR-ARNT association (**Fig. 5b-d**), demonstrating the sensitivity of the assays to competitive binding of the common partner. Given that AHR participates in both canonical and non-canonical signaling pathways^52,53^, and shares ARNT with multiple transcriptional complexes^37,54^, these assays enable quantitative analysis of pathway crosstalk at the protein interaction level. NanoBiT has been previously employed to monitor heterodimerization between HIF-1α and ARNT, mainly to assess effects of drugs stabilizing or inhibiting HIF-1α in the signaling complex^55,56^. Relative to HIF, adoption of either NanoBiT or BiFC techniques to AHR has the advantage of AHR activation being triggered by added ligands.

In summary, NanoBiT and BiFC provide sensitive and versatile platforms for direct measurement of AHR activation through protein-protein interaction analysis. By focusing on interaction events upstream of transcriptional outputs, these assays afford hitherto unavailable approaches while reducing ambiguity arising from context-dependent gene regulation. Beyond monitoring AHR-ARNT heterodimerization, they enable interrogation of early steps in AHR activation and can facilitate analysis of crosstalk between AHR and other pathways such as HIF signaling. Being ligand dependent, our assays may prove valuable for screening and activity profiling of AHR-active compounds while also enabling their detection in complex biological or environmental samples. More broadly, the modular design of these protein-based assays is adaptable to additional transcription factor networks, thereby extending protein-centered interrogation of transcription factor activity beyond AHR biology.

## Material & Methods

To support reproducibility of the experimental procedures described in this study, detailed information on all reagents, tools and sequences, including ordering information, is provided in **Supplementary Data 1**.

### Cell culture and preparation for experimental procedures

Chinese hamster ovary (CHO), LN-229 glioblastoma and human embryonic kidney (HEK293T) cells were purchased from ATCC and cultured at 37 °C in a humidified incubator with 5% CO_2_. CHO cells were used for all NanoBiT experiments, whereas LN-229 and HEK293T cells were used for BiFC, high resolution microscopy, XRE-Luciferase reporter assay and gene expression analyses via qRT-PCR.

CHO cells were cultured in high-glucose DMEM + GlutaMax^TM^ (Gibco, Thermo Fisher Scientific) supplemented with 10% heat-inactivated fetal bovine serum (FBS) and 1% penicillin-streptomycin (both Sigma-Aldrich). LN-229 and HEK293T cells were maintained in high-glucose DMEM (4.5 g/L glucose; Gibco, Thermo Fisher Scientific) supplemented with 10% FBS (Thermo Fisher Scientific), 1 mM sodium pyruvate, 2 mM L-glutamine, 100 U/mL penicillin and 100 µg/mL streptomycin (all Gibco, Thermo Fisher Scientific).

Mycoplasma contamination was routinely monitored using a commercial detection kit (Minerva Biolabs). At 90-100% confluency, cells were washed once with phosphate-buffered saline (PBS) and detached using trypsin-EDTA (Gibco, Thermo Fisher Scientific). Cells were resuspended in complete medium and passaged as required. Cells between passages 5 and 25 were used to minimize experimental variability.

### Design of protein expression vectors

Protein-protein interactions of three proteins (coding sequences (CDSs) in brackets) were investigated: aryl hydrocarbon receptor (AHR) (CCDS-ID: 5366.1), aryl hydrocarbon receptor nuclear translocator (ARNT) (CCDS-ID: 970.1) and heat shock protein 90 alpha family class b member 1 (HSP90AB1, further referred to as HSP90) (CCDS-ID: 4909.1).

Constructs encoding AHR fused to NanoLuc luciferase fragments comprised the full-length human AHR protein (M1-L848; clone RC2099832, OriGene; RefSeq NM_001621) with an N-terminal Myc epitope tag. The NanoLuc fragment was separated from AHR by a 22-amino acid linker (**Supplementary Data 1**). ARNT constructs encoded a C-terminally truncated form of human ARNT (M1-Q503) containing the bHLH and both PAS domains required for heterodimerization with AHR. ARNT was fused to NanoLuc fragments via an 18-amino acid linker and carried an N-terminal Flag epitope tag (**Supplementary Data 1**). AHR and ARNT constructs with N-terminal fusions to either the large (LgBiT) or small (SmBiT) NanoLuc fragments were generated in the pBiT1.1-N [TK/LgBiT] or pBiT2.1-N [TK/SmBiT] vectors (Promega), respectively. These vectors were modified to introduce additional restriction sites and N-terminal Myc or Flag epitope tags (**Supplementary Data 1**).

Site-directed mutagenesis of AHR was performed to introduce the Y322A substitution by PCR-based mutagenesis using complementary primers (forward: 5′-GCACGAGAGGCTCAGGT<u>GC</u>TCAGTTTATTCATGCAGCTG-3′; reverse: 5′-CAGCTGCATGAATAAACTGA<u>GC</u>ACCTGAGCCTCTCGTGC-3′), replacing the corresponding wild-type nucleotides (mutated residues underlined). Mutant constructs were cloned into the same NanoBiT vectors as the wild-type AHR constructs.

A construct encoding full-length human HSP90 (UniProt P08238) in the pFN21A vector was obtained from Promega (FHC01815). HSP90 was fused at the N terminus to the LgBiT NanoLuc fragment by subcloning into the corresponding pBiT vectors (**Supplementary Data 1**).

All constructs were verified by sequencing prior to use. Protein-protein interactions were analyzed via NanoBiT using the following combinations: LgBiT-linker-Myc-AHR (LgAHR) with SmBiT-linker-Flag-ARNT(1–503) (SmARNT), and LgBiT-linker-Flag-HSP90 (LgHSP90) with SmBiT-linker-Myc-AHR (SmAHR).

For BiFC experiments, the CDS of human HSP90 (NM_001271969) was obtained as Gateway®-compatible entry clone in pDONR223 from the clone repository of the DKFZ Genomics and Proteomics Core Facility (GPCF). To generate AHR and ARNT expression vectors, AHR (NM_001621) and ARNT (NM_001668) cDNA was amplified by PCR and cloned into the Gateway® entry vector pDONR201, obtained from the GPCF, using primers annealing to the start or end of the AHR and ARNT CDSs. AttB recombination sites were introduced using a two-step PCR procedure. In the first step, hybrid primers containing partial AttB sequences and gene-specific regions were used, followed by amplification with primers spanning the complete AttB sites (for primer sequences see **Supplementary Data 1**). After sequence verification, the resulting pDONR201-AHR and pDONR201-ARNT entry clones were used for subsequent Gateway® cloning.

Gateway® destination vectors pGW-MYC-LC151 and pGW-HA-LN151, which encode C-terminal MYC or HA epitope tags fused to the respective bimolecular fluorescence complementation (BiFC) fragments have been described previously^57,58^. These vectors were additionally used as negative controls to assess the specificity of protein-protein interactions and served as entry clones for generating the final expression constructs via Gateway® LR recombination: pGW-AHR-HA-LN151, pGW-ARNT-MYC-LC151 and pGW-HSP90-MYC-LC151. In all constructs, the epitope tags and BiFC fragments were fused to the C terminus of the respective protein coding sequences.

BiFC and respective control plasmids were propagated in *Escherichia coli* DB3.1 or Stbl3 strains and purified using the QIAprep Spin Miniprep Kit (Qiagen) or the NucleoBond Xtra Midi Kit (Macherey-Nagel).

### NanoBiT luciferase assay

NanoBiT technology (Promega) was used to monitor protein-protein interactions in live cells as previously described^22^ and NanoBiT assays were performed as previously reported^59^. Here, we used NanoBiT to monitor assembly of AHR-ARNT and dissociation of AHR-HSP90 complexes. CHO cells were seeded at semiconfluent density (∼2 × 10L cells/well) in flat-bottom, white, solid 96-well plates (Corning). 24 hours (h) after seeding, cells were transiently transfected with 200 ng total plasmid DNA per well at a 1:1 ratio of the respective two vectors using FuGENE® HD (Promega), following the manufacturer’s instructions.

After an additional 24 h, cells were equilibrated for 10 min at room temperature (RT) in serum-free DMEM buffered with 20 mM HEPES (pH 7.2; HDMEM). Nano-Glo® Live Cell Substrate (Promega) was then added, and baseline luminescence was recorded for 20 min using an Orion II microplate luminometer (Berthold Technologies).

AHR ligands, including 6-formylindolo[3,2-b]carbazole (FICZ) and indirubin (both Sigma-Aldrich), as well as the AHR antagonist KYN-101 (MedChemExpress), were dissolved in DMSO to generate stock solutions of 10 mM (FICZ and KYN-101) or 5 mM (indirubin). DMSO at a final concentration of 0.05% was used as vehicle control. FICZ and indirubin were typically used at a final concentration of 5 nM, unless stated otherwise. Ligands or vehicle controls were added manually from 10x stocks in HDMEM using fresh pipette tips for each replicate, and luminescence was recorded for an additional 30 min. Baseline luminescence values varied between wells, most likely reflecting differences in transfection efficiency. To account for this variability, NanoBiT signals were normalized to the baseline luminescence of each well.

### BiFC assay

Bimolecular fluorescence complementation (BiFC) was used to monitor assembly of AHR-ARNT complexes and dissociation of AHR-HSP90 complexes in live cells. Briefly, non-fluorescent N-and C-terminal fragments of the far-red fluorescent protein mLumin, a member of the mKate protein family^60^, were fused to the coding sequences of AHR, ARNT and HSP90, as described above.

LN-229 and HEK293T cells were counted using an automated cell counter (Countess 3, Thermo Fisher Scientific). Approximately 8.0 × 10L LN-229 cells or 1.6 × 10L HEK293T cells were seeded in 1 mL DMEM per well in 24-well plates (µ-Plate 24 Well; Ibidi). 20 h after seeding, cells were transiently transfected with 750 ng total plasmid DNA per well at a 1:1 ratio of the respective BiFC vectors using the Lipofectamine™ 3000 Transfection Reagent Kit (Thermo Fisher Scientific). Transfections were performed using 2 µL P3000 reagent per µg DNA and 1 µL Lipofectamine 3000 per well, according to the manufacturer’s instructions, in a final volume of 50 µL Opti-MEM (Gibco), which was added to the pre-existing culture medium. After 4-5 h, the transfection mixture was removed and replaced with fresh DMEM containing the indicated AHR ligands, stool samples or carrier-controls.

AHR ligands, including 6-formylindolo[3,2-b]carbazole (FICZ; MedChemExpress), indirubin (Sigma-Aldrich), kynurenic acid (KynA; Sigma-Aldrich) and indole-3-pyruvate (I3P; Sigma-Aldrich), as well as the AHR antagonist KYN-101 (MedChemExpress), were dissolved in DMSO to generate stock solutions of 50 mM (KynA and I3P), 10 mM (KYN-101) or 1 mM (FICZ and indirubin). FICZ and indirubin were typically used at a final concentration of 100 nM, KynA at 100 µM and I3P at 25 µM, unless stated otherwise. Dimethyloxaloylglycine (DMOG), a hypoxia-mimetic compound, was dissolved in DMSO to prepare a 2 M stock solution and used at a final concentration of 2 mM. Ligands, vehicle controls or stool samples were prepared in fresh DMEM and applied to the cells after removal of the transfection medium. Stool samples are further described in the section “human stool samples”. Cells were imaged approximately 22 h after ligand treatment.

### Microscopy

Microscopy was performed using a Leica MICA Microhub. Prior to analyzing protein-protein interactions, appropriate control experiments were conducted to assess potential nonspecific interference in the fluorescence channel. For this purpose, mock-transfected conditions were included, along with constructs encoding the respective BiFC fragments lacking the coding sequences (CDS) of AHR, ARNT, or HSP90. Subsequently, AHR-ARNT and AHR-HSP90 interactions were visualized by BiFC.

Live, unfixed LN-229 and HEK293T cells cultured in µ-Plate 24-well plates were imaged. Plates feature a flat, optically clear polymer coverslip bottom and black walls to minimize cross-well signal interference. All images used for quantification were acquired in air (refractive index n = 1.000) using a 10× HC PL FLUOTAR dry objective (NA 0.32, working distance 11.2 mm). For each well, approximately 30 non-overlapping fields of view were acquired at predefined positions around the central area of the well to avoid edge-related artifacts. At each position, one transmitted light (TL) image and one fluorescence (FL) image were recorded. TL images were acquired in integrated modulation contrast (IMC) mode (offset 0, light intensity 128, exposure time 40 ms, gain 1.0). FL images were acquired in widefield mode using the Leica mKate2 excitation profile (excitation 589 nm), achieved using LED 555 illumination. For each experiment, LED intensity and exposure time were optimized at the beginning of image acquisition and subsequently kept constant across all wells within that experiment. Exposure times ranged from 100-250 ms, with a gain of 5.0. Emission was detected using the predefined mKate2 detection channel (center wavelength 633 nm) with a 5 MP CMOS camera. Images were acquired at a resolution of 2432 × 2032 pixels, corresponding to a field of view of 1.398 × 1.168 mm and a pixel size of 575 nm.

Image analysis was performed using a custom-made ImageJ macro (**Supplementary Data 2**). The macro quantifies both the total cell-covered area per image and the area of BiFC-derived fluorescence above a defined threshold following background subtraction, which was kept constant across all biological experiments. In detail, the fluorescence (FL) channel was processed by applying background subtraction (rolling ball radius = 50), followed by Gaussian blurring and segmentation using a fixed threshold (150-65535). Particles with an area greater than 20 pixels were included in the analysis. For the transmitted light (TL) channel, a variance filter (radius = 10) was applied to enhance structural features. Subsequently, contrast was enhanced, a lookup table (LUT) was applied, and the image was converted to 8-bit. Segmentation was then performed using a fixed threshold (30-255), and particles larger than 800 pixels were considered for the final analysis. Images containing artifacts such as debris, uneven illumination or interference from plate texture were excluded before analysis.

Images were processed for publication using the open-source Open Microscopy Environment (OMERO) platform, specifically OMERO.insight and OMERO.figure. Representative microscopy images were prepared for publication using OMERO.figure. Scale bars (500 or 100 µm, as indicated in the figure legends) were automatically generated from image metadata. The BiFC signal, originally detected in the red spectrum (mLumin), was pseudocolored in magenta to improve visual clarity.

### High resolution microscopy

BiFC-transfected cells were prepared for monitoring subcellular localization of protein-protein interactions using high-resolution microscopy. For these nuclear staining experiments, the same cell culture plates were used as in the corresponding BiFC experiments, with cells seeded, transfected, and treated identically. Then culture medium was carefully removed and cells were directly fixed with 4% formaldehyde for 11 min at RT. The fixation solution was prepared in advance by dissolving 4% paraformaldehyde (PFA; Sigma-Aldrich) in PBS under heating. Following fixation, cells were washed with PBS and permeabilized using 0.2% Triton X-100 (Sigma-Aldrich) in PBS for 10 min. After an additional washing step, cells were incubated for 10 min in PBS containing 1 µg/mL 4′,6-diamidino-2-phenylindole (DAPI; SERVA) for nuclear staining. Subsequently, the staining solution was removed, cells were washed once with PBS, and fresh PBS was added prior to imaging.

Imaging was performed using a 63× HC PL APO CS2 oil immersion objective (NA 1.4, WD 140 µm). Type F immersion oil (Thermo Fisher Scientific) was applied. Transmitted light (TL) and fluorescence (FL) images for BiFC were acquired as described above. DAPI fluorescence was recorded using the integrated “DAPI (dsDNA bound)” channel with LED365 illumination. Fresh immersion oil was applied for each imaging position. Imaging with the 63× objective resulted in a field of view of 221.9 µm × 185.4 µm, corresponding to a pixel size of 91 nm, while maintaining consistent image resolution settings across experiments.

Representative images were prepared using OMERO.figure. Scale bars (25 µm, as indicated in the figure legends) were automatically generated from image metadata. The BiFC signal, originally detected in the red spectrum (mLumin), was pseudocolored in magenta, while the DAPI signal, originally detected in the blue spectrum, was pseudocolored in white to enhance visual clarity.

### Gene expression analysis via qRT-PCR

RNA was isolated immediately after microscopy. For non-transfected conditions, cells were seeded and treated in parallel under identical conditions.

Total RNA was isolated using the RNeasy Mini Kit (QIAGEN) according to the manufacturer’s instructions. Cells were lysed in RLT buffer supplemented with 1% β-mercaptoethanol (Sigma-Aldrich), and lysates were subjected to on-column DNase (QIAGEN) digestion. RNA was eluted in RNase-free water, and concentrations were determined using a FastGene NanoSpec Photometer (NIPPON Genetics).

cDNA synthesis was performed using the High-Capacity cDNA Reverse Transcription Kit (Thermo Fisher Scientific) according to the manufacturer’s instructions. Briefly, 1 µg of total RNA was combined with reaction buffer, random hexamer primers, reverse transcriptase, and nuclease-free water. Reverse transcription was carried out in a T3000 Thermocycler (Analytik Jena) under the following conditions: 10 min at 25 °C, 120 min at 37 °C, and 5 min at 85 °C.

For gene expression analysis, cDNA was combined with SYBR Green Master Mix (Thermo Fisher Scientific), gene-specific forward and reverse primers, and nuclease-free water, and loaded in technical triplicates into a 96-well PCR plate (Steinbrenner) in a total reaction volume of 15 µL per well. Primers targeting the housekeeping gene *18S rRNA* were included on each plate as an internal reference (for primer sequences see **Supplementary Data 1**).

Plate layout and cycling conditions were designed using Design and Analysis 2 software and run on a QuantStudio 3 Real-Time PCR System (both Thermo Fisher Scientific). qPCR was performed under the following conditions: 10 min at 95 °C, followed by 40 cycles of 15 s at 95 °C and 60 s at 60 °C. Melting curve analysis was conducted with 15 s at 95 °C, 60 s at 60 °C, and 15 s at 95 °C. Plates were sealed with adhesive film (Thermo Fisher Scientific) prior to machine loading.

Data were analyzed using Design and Analysis 2 software, which automatically determined cycle threshold (Ct) values for each well. For each sample, Ct values were calculated as the mean of technical triplicates. If the standard deviation exceeded 0.25, the outlier was excluded, and the mean of the two closest Ct values was used. Relative gene expression was calculated using the comparative Ct (ΔΔCt) method with *18S rRNA* as the internal reference, as previously described^61^. Fold change (FC) was determined as:

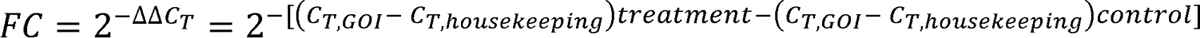

### Immunoblot analysis

Expression of NanoBiT-tagged and BiFC-tagged proteins was confirmed by immunoblotting.

For NanoBiT-tagged proteins, FugeneHD-transfected CHO cells (3 µg total plasmid DNA/well at a 1:1 ratio of the respective constructs) in 6-well plates were washed in ice-cold PBS and lysed in Passive Lysis Buffer (Promega). Proteins were separated by SDS-PAGE on 8% polyacrylamide gels under reducing conditions using LDS sample buffer (Promega) supplemented with 1.25% β-mercaptoethanol (Serva). Proteins were transferred to nitrocellulose membranes (Cytiva) and blocked in buffer containing 0.1% Tween 20 (AppliChem), 50 mM Trizma base (pH 7.8, Sigma-Aldrich), 0.2 M NaCl (Serva), 1 mM EDTA (Sigma-Aldrich), with 5% nonfat dried milk (AppliChem) and 5% bovine albumin, fraction V (Serva).

Wild-type and mutant AHR proteins, all carrying an N-terminal Myc epitope, were detected using the monoclonal anti-Myc antibody 9E10 (1:2000 in blocking buffer; Thermo Fisher Scientific), followed by incubation with HRP-conjugated secondary antibodies and chemiluminescent detection with SuperSignal West Femto Maximum Sensitivity Substrate (Thermo Fisher Scientific) using an Uvitec Alliance Q9 imaging system (UVItec Cambridge). Flag-tagged ARNT and HSP90 proteins were detected using a monoclonal anti-Flag antibody (MA1-91878, 1:2000; Thermo Fisher Scientific). Ribosomal protein S6 (S6RP) was used as a loading control (1:2000; Cell Signaling Technology).

For immunoblotting of BiFC-tagged proteins, approximately ∼8 × 10L HEK293T cells were seeded per well in 6-well plates (Greiner Bio-One). The following day, cells were transiently transfected with 5 µg total plasmid DNA per well at a 1:1 ratio of the respective BiFC constructs using Lipofectamine 3000. Transfections were performed with 2 µL P3000 reagent per µg DNA and 4 µL Lipofectamine 3000 per well in 250 µL Opti-MEM, added directly to the culture medium according to the manufacturer’s instructions. Mock-transfected controls received transfection reagents without plasmid DNA. After 4-5 h, the transfection mixture was replaced with fresh DMEM. Cells were harvested for protein isolation after an additional 24 h. Protein isolation and immunoblotting were performed as previously described^58^. All steps were carried out on ice unless otherwise stated to minimize protein degradation. For protein isolation, culture medium was removed and cells were washed once with ice-cold PBS. Cells were lysed in RIPA buffer (sterile DPBS supplemented with 1% IGEPAL/NP-40 (Sigma-Aldrich), 0.5% sodium deoxycholate (AppliChem), and 0.1% sodium dodecyl sulfate (SDS) (Carl Roth); sterile-filtered and stored at 4 °C). Immediately prior to use, the buffer was supplemented with 1% phosphatase inhibitor cocktails 2 and 3 (Sigma-Aldrich) and 4% complete protease inhibitor (Roche). Cells were scraped, and lysates were incubated on ice for 5 min, followed by centrifugation at 13,000 × g for 10 min at 4 °C. The supernatant was collected and protein concentration was determined using Protein Assay Dye Reagent Concentrate (Bio-Rad) by measuring absorbance at 595 nm with a spectrophotometer (Amersham Biosciences). Protein concentrations were used to normalize samples prior to the addition of 5x Laemmli buffer (10% glycerol (Sigma-Aldrich), 1% β-mercaptoethanol, 1.7% SDS, 62.5 mM Trizma base (Sigma-Aldrich), pH 6.8, bromophenol blue (Sigma-Aldrich), and dH₂O), which was preheated to 95 °C.

For SDS-PAGE, self-cast 10% polyacrylamide gels were prepared as previously described^58^. Gels were mounted in an electrophoresis chamber and filled with running buffer (25 mM Trizma base, 0.2 M glycine (Sigma-Aldrich), 0.1% SDS). Protein samples were denatured at 95 °C for 5 min and loaded (11 µL per lane) on polyacrylamide gels. Protein ladders (Precision Plus Protein™ Dual Color Standards (Bio-Rad); and PageRuler™ Plus Prestained Protein Ladder, 10-250 kDa (Thermo Fisher Scientific)) were loaded (4 µL each) as reference. Electrophoresis was performed at 80 V for 10-15 min, followed by 120 V for 1.5-2 h, depending on protein size.

Proteins were transferred onto methanol-activated PVDF membranes (Merck Millipore) using a wet transfer system. Membranes were assembled with Whatman filter papers (Cytiva) and sponge pads in transfer buffer (50 mM Trizma base, 0.1 M glycine, 0.01% SDS, 10% methanol (Thermo Fisher Scientific), pH 8.3) and transferred at 45 V for 2 h with cooling. Membranes were briefly rinsed in methanol, dried, labeled, cut, methanol-reactivated and blocked in 2.5% BSA (Carl Roth) in TBS-T (0.15 M NaCl (Sigma-Aldrich), 60 mM Trizma base, 3 mM KCl (Sigma-Aldrich), 0.1% Tween-20 (Sigma-Aldrich), pH 7.4) for 1 h at RT with gentle agitation.

Membranes were incubated overnight at 4 °C with primary antibodies diluted in blocking buffer (5% BSA in TBS-T supplemented with 0.1% sodium azide (AppliChem)) according to the manufacturer’s instructions. Membranes were washed three times briefly, followed by three additional washes of 10 min each in TBS-T. Subsequently, membranes were incubated with the appropriate HRP-conjugated secondary antibodies, freshly diluted 1:4000 in TBS-T, for 2 h at RT. After washing, signals were detected using SuperSignal™ West Femto Maximum Sensitivity Substrate (Thermo Fisher Scientific) and imaged with a ChemiDoc MP Imaging System (Bio-Rad).

Membranes were re-probed by repeating the same procedure to detect loading controls. Primary antibodies included anti-HA High Affinity (1:1000; Roche), anti-Myc (1:1000; Cell Signaling Technology), and anti-Tubulin (1:10000; TUBA1B; Abcam). Corresponding HRP-conjugated secondary antibodies (goat anti-rat, goat anti-mouse, and goat anti-rabbit; Thermo Fisher Scientific) were used.

### XRE-Luciferase Reporter Assay

Approximately 1.5 × 10L LN-229 cells were seeded per well in 96-well Nunclon Delta-treated white flat-bottom plates (Thermo Fisher Scientific) in 100 µL DMEM. The following day, cells were co-transfected with 100 ng total plasmid DNA per well consisting of pGL4.43 [luc2P/XRE/Hygro] and pGL4.74 [hRluc/TK] (Promega) at a 10:1 ratio using Lipofectamine 3000 (Thermo Fisher Scientific). Transfections were performed with 2 µL P3000 reagent per µg DNA and 0.25 µL Lipofectamine 3000 per well in 10 µL Opti-MEM, added directly to the culture medium. After 18 h, the transfection mixture was replaced with fresh DMEM containing the indicated AHR ligands (75 µL per well).

After 24 h, luciferase activity was measured using the Dual-Glo® Luciferase Assay System (Promega) according to the manufacturer’s instructions. Plates were equilibrated to RT for 15 min prior to reagent addition. Dual-Glo Luciferase Reagent (75 µL per well) was added, incubated for 10 min, and firefly luciferase activity was measured using a CLARIOstar plate reader (BMG Labtech) (top optics, emission 580 nm, gain 3600, orbital averaging scan, 4 mm diameter, 25 intervals, 2.47 s interval time). Subsequently, 75 µL Dual-Glo Stop & Glo Reagent was added, incubated for 10 min, and Renilla luciferase activity was measured under identical settings (emission 480 nm). Focal height was determined automatically and kept constant across the plate. Measurements were performed in technical triplicates. Firefly luciferase activity was normalized to Renilla luciferase activity for each well, and values were expressed relative to the vehicle control (DMSO).

### Human stool samples

Human stool samples were obtained from three individual healthy volunteers under the ethics board approval S-496/2014.

Stool samples were diluted 1:15 in amino extract buffer (Immundiagnostik AG) and PBS and allowed to settle to enable particulate matter to sediment. The resulting supernatant was then centrifuged at 10,500 × g for 60 min. Following centrifugation, the supernatant was subjected to microfiltration using a Minisart® syringe filter (0.20 µm; Sartorius) and stored at −20 °C until further use. For BiFC experiments, cells were treated with processed stool samples, or PBS as control, at dilutions of ≈1:6, corresponding to a final stool concentration of ≈1:100.

### Statistical analysis

Statistical analyses and data visualization were performed using GraphPad Prism (version 11.0.0). Technical replicates are presented as mean ± SD (standard deviation), whereas biological replicates are reported as mean ± SEM (standard error of the mean). Statistical analyses were performed only when at least three independent biological replicates were available.

BiFC experiments were analyzed by quantifying the percentage of fluorescence-positive area (**Extended Data Fig. 1**). Individual values were normalized to the mean of the carrier control (DMSO) to calculate fold changes. For NanoBiT analyses, relative luciferase activity for each well was determined as a ratio between the measured luminescence and the respective baseline value (substrate only) at the time of ligand addition (t = 0 min, set to 1) and displayed as relative fold change. For NanoBiT dose-response measurements (**Fig. 2a, f**; **Fig. 3a, f**), relative luminescence [%] was determined by normalizing all values to the maximum response achieved for the highest ligand concentration used, which was set to 100%. The baseline at the timepoint of ligand addition (t = 0 min) was set to 0. EC₅₀ values were determined by nonlinear regression analysis using fold-change data (BiFC experiments) or area-under-the-curve (AUC) values (NanoBiT). For NanoBiT analyses, the vehicle control (“0 concentration”) was included in the regression as indicated in the figure legends (**Extended Data Fig. 3a, b**).

For hypothesis testing, fold-change values were log-transformed to approximate a normal distribution. One-sample *t*-test was performed against a theoretical mean of 0. When multiple conditions were compared to the respective control, resulting *p*-values were adjusted for multiple comparisons using the Holm-Šidák method.

The number of biological replicates and the exact adjusted *p*-values are provided in the corresponding figure legends as well as in **Supplementary Data 3**. Statistical significance is indicated as follows: *p* < 0.05 (\**), p < 0.01 (**), p < 0.001 (\*\*\**), and *p* < 0.0001 (****); ns, not significant (*p* > 0.05).

## Data availability

Requests for resources and reagents should be directed to the corresponding authors Mirja Tamara Prentzell, Christiane A. Opitz and Sarka Tumova.

Source data underlying the figures will be provided with this paper. Raw microscopy image datasets are available from Tamara Prentzell upon reasonable request.

## Acknowledgements

The authors thank Lara Elea Eckhardt, Martin Herdt, Lukas Voos, Ivana Karabogdan and Soumya Mohapatra for experimental support as well as Damir Krunic for help with the ImageJ macro. The authors acknowledge the entire Metabolic Crosstalk in Cancer lab for fruitful discussions and valuable feedback.

CAO acknowledges support from the German Research Foundation (SFB1389 *UNITE-Glioblastoma*; project no. 404521405) and the European Research Council (ERC; grant agreement no. 101045257, *CancAHR*). MTP and CAO further acknowledge funding from the Central Innovation Program for SMEs (ZIM) of the former Federal Ministry for Economic Affairs and Energy (BMWI, grant no. ZF4101406SK8, *Universeller Aryl Hydrocarbon Rezeptor Assay*), and from the Federal Ministry for Economic Affairs and Climate Action (BMWK; grant no. KK5448002AJ2, *Healthy Gut*). KK also acknowledges funding from the BMWI (grant no. ZF4647301SK8, *Universeller Aryl Hydrocarbon Rezeptor Assay*) and the BMWK (grant no. KK5522101AJ2, *Healthy Gut*). NZ was supported by a U.S. Fulbright Research Scholarship (2025-2026). **Fig. 1a, Fig. 1b**, **Fig. 5a**, **Fig. 5e**, **Extended Data Fig. 1a, Extended Data Fig. 1b** and **Extended Data Fig. 1c** were created using BioRender with the following agreement numbers: **VY29TQSRFU, DW29TQSX68, HQ29TQT3Q2, UV29TQTBG2, JV29TQQWBL, MM29TQS13I, AU29TQS9AX**.

## Author contributions

**Tim Kühn:** Formal analysis; Investigation; Methodology; Visualization; Writing – review and editing. **Sarka Tumova:** Conceptualization; Formal analysis; Investigation; Methodology; Visualization; Writing – review and editing. **Nicholas Zacharewski:** Investigation. **Paul Johannes Averdung:** Investigation. **Bianca Berdel:** Investigation. **Karl-Heinz Kellner:** Resources. **Stefan Pusch:** Resources. **Marek Jindra:** Conceptualization; Funding acquisition; Methodology; Writing – review and editing. **Christiane A. Opitz:** Conceptualization; Funding acquisition; Methodology; Supervision; Writing – review and editing. **Mirja Tamara Prentzell:** Conceptualization; Funding acquisition; Methodology; Project administration; Supervision; Writing – original draft; Writing – review and editing. All authors have read and agreed to the manuscript.

## Declaration of generative AI and AI-assisted technologies in the writing process

In preparing this work, the authors used ChatGPT, an AI language model developed by OpenAI, to enhance the readability and clarity of the manuscript without altering its scientific meaning or integrity. Following the use of this tool, the authors carefully reviewed and edited the content to ensure accuracy and originality, and they accept full responsibility for the published article.

## Supplementary Data

**Supplementary Data 1:** Reagents and Tools Table.

**Supplementary Data 2:** ImageJ macro for automated quantification of BiFC-mediated fluorescence.

**Supplementary Data 3**: *p*-value overview.

**Supplementary Data 4:** Vector maps of used NanoBiT and BiFC plasmids.

## Extended Data Figure Legends

**Extended Data Fig. 1.**
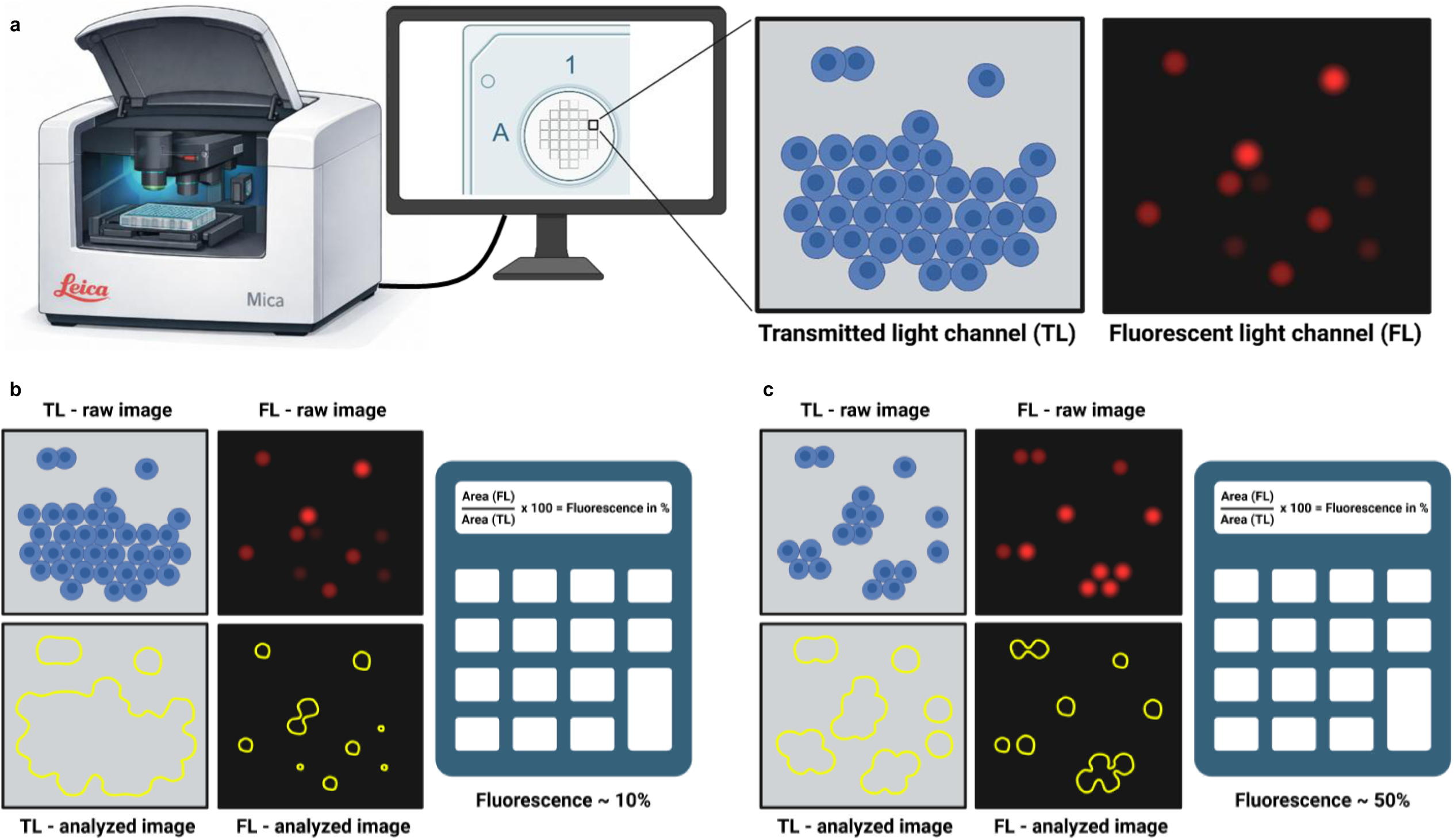
BiFC image analysis pipeline. **a,** Overview of the BiFC imaging workflow. Cells were seeded in 24-well plates, transfected with BiFC constructs and imaged using the Leica MICA Microhub. Up to 30 independent, non-overlapping fields of view were acquired per well. For each field of view, transmitted light (TL) and fluorescence (FL) images were recorded. Image created with BioRender and ChatGPT. **b,c,** Representative TL and FL images illustrating quantitative image analysis using a custom ImageJ macro (see Supplementary Data 2). For each image, the fluorescent area (Area FL) was normalized to the corresponding cell-covered area derived from the TL image (Area TL), yielding fluorescence expressed as a percentage per image. Panel **b** shows an example with 10% normalized fluorescence, whereas panel **c** shows an image with 50% normalized fluorescence. These values indicating the normalized fluorescence-positive area are then further normalized and expressed as fold change relative to DMSO. Image created with BioRender.

**Extended Data Fig. 2.**
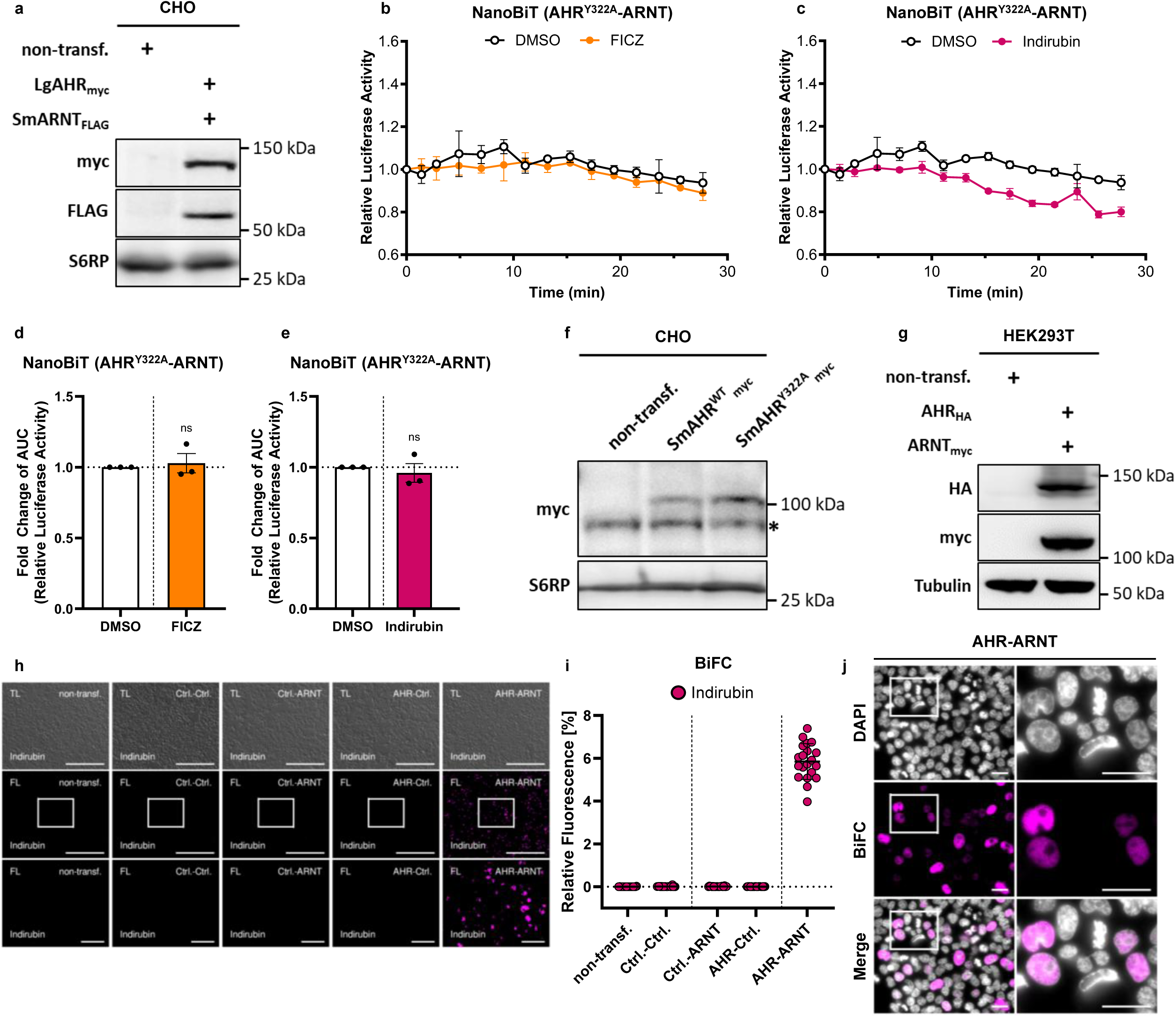
AHR-ARNT interaction occurs in the nucleus and mutagenesis of the ligand binding pocket prevents AHR-ARNT heterodimerization. **a**, Representative immunoblot of CHO cells either non-transfected or transfected with NanoBiT constructs encoding AHR fused to LgBiT (LgAHR_myc_) and ARNT fused to SmBiT (SmARNT_FLAG_). Expression of the NanoBiT fusion proteins was confirmed by detection of the myc- and FLAG-epitope tags. Ribosomal protein S6 (S6RP) was used as a loading control. Representative of n = 3 independent biological replicates. **b, c**, Representative NanoBiT time-course of AHR-ARNT protein-protein interaction using a mutated AHR construct (AHR^Y322A^) in CHO cells, showing relative luciferase activity upon treatment with 5 nM FICZ (b) or indirubin (c) compared to DMSO control, corresponding to AHR wildtype (AHR^WT^) as shown in Fig. 1c. Data are shown as mean ± SD of two technical replicates. **d, e**, Quantification of NanoBiT luminescence data for AHR^Y322A^ (examples shown in b, c). The area under the curve (AUC) was calculated for each condition and expressed as fold change relative to DMSO (set to 1). Data are mean ± SEM of n = 3 independent biological replicates. Statistical significance was assessed using one-sample *t*-test on log-transformed fold-change values. d, *p* = 0.75 (ns). e, *p* = 0.56 (ns). **f**, Representative immunoblot of CHO cells either non-transfected or transfected with NanoBiT constructs encoding AHR^WT^ fused to SmBiT (SmAHR^WT^) or AHR^Y322A^ fused to SmBiT (SmAHR^Y322A^_myc_). Expression of the NanoBiT fusion proteins was confirmed by detection of the myc epitope tags. Ribosomal protein S6 (S6RP) was used as a loading control. * denotes nonspecific background band. Representative of n = 2 independent biological replicates. **g**, Representative immunoblot of HEK293T cells either non-transfected or transfected with BiFC constructs encoding AHR and ARNT fused to complementary fluorescent protein fragments. Expression of the BiFC fusion proteins was confirmed by detection of the HA- and myc-tagged AHR and ARNT, respectively. Tubulin was used as a loading control. Representative of n = 5 independent biological replicates. **h**, Representative BiFC microscopy images including negative BiFC controls following treatment with 100 nM indirubin. Top, transmitted light (TL); middle, fluorescence (FL); bottom, magnified views of the boxed regions in the FL images. Fluorescence signals are pseudocolored in magenta for visualization. Scale bars, 500 µm (top and middle panels) and 100 µm (bottom panel). **i**, Representative BiFC experiment in HEK293T cells showing normalized fluorescence-positive area upon treatment with 100 nM indirubin, expressed in %, with representative images shown in h. Cells were either non-transfected, transfected with control BiFC vectors lacking the coding sequences of AHR and ARNT, but containing the corresponding fluorescent fragments, or co-transfected with AHR and ARNT BiFC constructs. Each dot represents a single, independent, non-overlapping field of view. Data are shown as mean ± SD of n ≥ 19 images per condition; n = 2 independent biological experiments. **j**, Representative BiFC microscopy images of AHR-ARNT protein-protein interaction, combined with nuclear DAPI staining in HEK293T cells. Top, DAPI channel; middle, BiFC channel; bottom, merged images. Enlarged views of the boxed regions in the left panels are shown on the right. DAPI signals are shown in white and BiFC signals are pseudocolored in magenta for visualization. Scale bars, 25 µm. Representative of n = 3 independent biological experiments.

**Extended Data Fig. 3.**
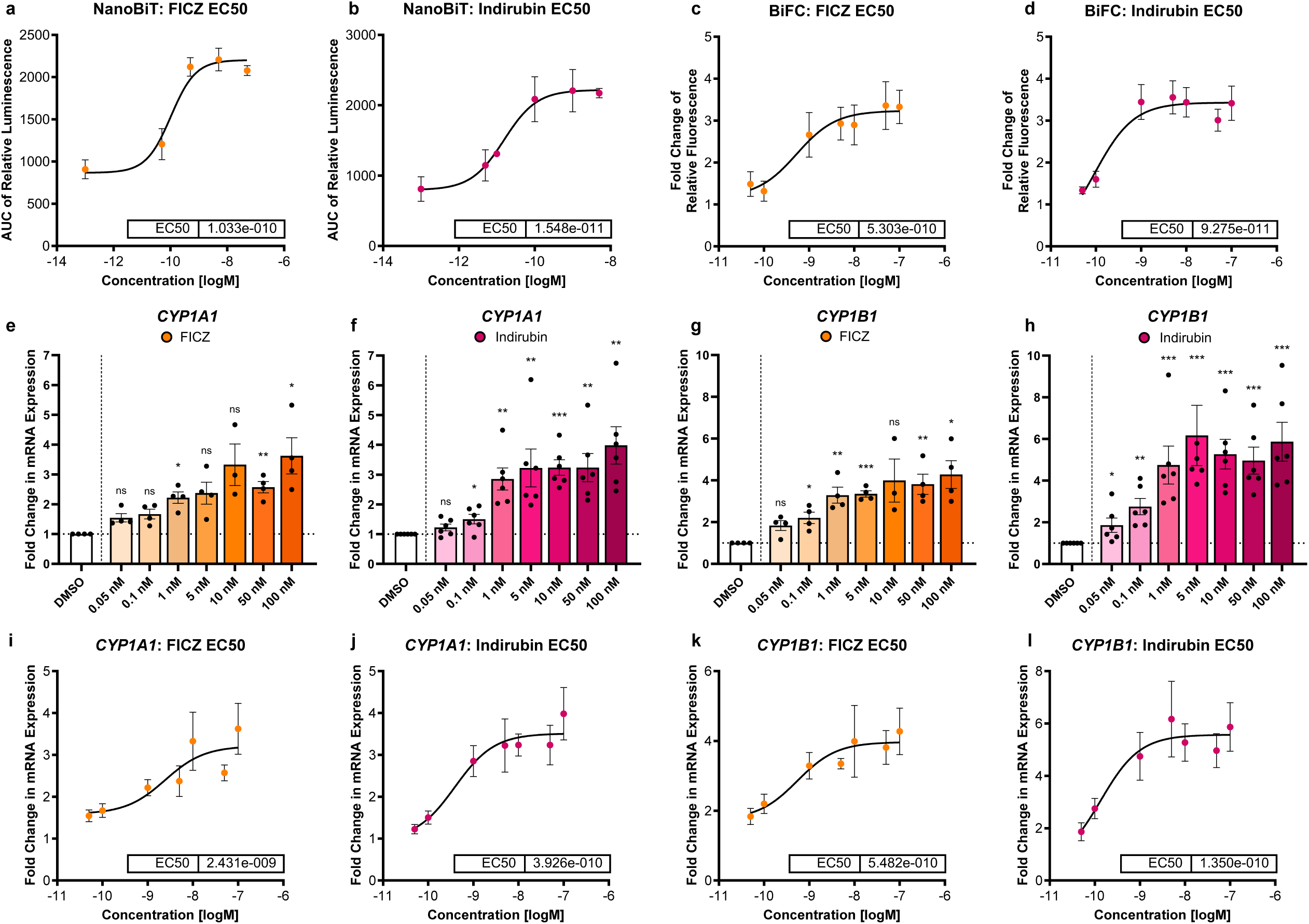
Concentration-dependent detection of AHR activation by NanoBiT, BiFC and transcriptional readouts. **a, b** The EC₅₀ for FICZ (**a**) and indirubin (**b**) were calculated by nonlinear regression of AUC values from the NanoBiT experiments shown in Fig. 2b and 2g using log-transformed concentrations. The vehicle control (“0 concentration”) was included in the regression. **a**, EC_50_ = 100 pM. **b**, EC_50_ = 15 pM. **c, d** The EC₅₀ for FICZ (**c**) and indirubin (**d**) were calculated by nonlinear regression of fold change values from the BiFC experiments shown in Fig. 2d and 2i using log-transformed concentrations. **c**, EC_50_ = 530 pM. **d**, EC_50_ = 93 pM. **e**, **f**, **g**, **h**, Quantification of *CYP1A1* (**e**, **f**) and *CYP1B1* (**g**, **h**) mRNA expression in HEK293T cells following FICZ (**e**, **g**) or indirubin (**f**, **h**) treatment, measured by qRT-PCR and expressed as fold change relative to DMSO (set to 1). RNA was extracted following BiFC imaging, corresponding to Fig. 2d and Fig. 2i, respectively. Data are mean ± SEM of n = 4 independent biological replicates for FICZ (except n = 3 for 10 nM FICZ) or n = 6 independent biological replicates for indirubin. Statistical significance was assessed using one-sample *t*-test on log-transformed fold-change values, followed by Holm-Šidák correction for multiple comparisons. **e**, *p* = 5.79 × 10⁻^2^ for 0.05 nM FICZ (ns); *p* = 5.79 × 10⁻^2^ for 0.1 nM FICZ (ns); *p* = 1.94 × 10⁻^2^ for 1 nM FICZ (*); *p* = 5.79 × 10⁻^2^ for 5 nM FICZ (ns); *p* = 5.79 × 10⁻^2^ for 10 nM FICZ (ns); *p* = 8.01 × 10⁻^3^ for 50 nM FICZ (**); *p* = 2.15 × 10⁻^2^ for 100 nM FICZ (*). **f**, *p* = 0.11 for 0.05 nM indirubin (ns); *p* = 4.56 × 10⁻^2^ for 0.1 nM indirubin (*); *p* = 1.86 × 10⁻^3^ for 1 nM indirubin (**); *p* = 4.03 × 10⁻^3^ for 5 nM indirubin (**); *p* = 1.73 × 10⁻^4^ for 10 nM indirubin (***); *p* = 1.86 × 10⁻^3^ for 50 nM indirubin (**); *p* = 1.69 × 10⁻^3^ for 100 nM indirubin (**). **g**, *p* = 5.65 × 10⁻^2^ for 0.05 nM FICZ (ns); *p* = 2.93 × 10⁻^2^ for 0.1 nM FICZ (*); *p* = 9.82 × 10⁻^3^ for 1 nM FICZ (**); *p* = 7.73 × 10⁻^4^ for 5 nM FICZ (***); *p* = 5.65 × 10⁻^2^ for 10 nM FICZ (ns); *p* = 9.82 × 10⁻^3^ for 50 nM FICZ (**); *p* = 1.07 × 10⁻^2^ for 100 nM FICZ (*). **h**, *p* = 2.37 × 10⁻^2^ for 0.05 nM indirubin (*); *p* = 1.85 × 10⁻^3^ for 0.1 nM indirubin (**); *p* = 8.89 × 10⁻^4^ for 1 nM indirubin (***); *p* = 8.89 × 10⁻^4^ for 5 nM indirubin (***); *p* = 3.76 × 10⁻^4^ for 10 nM indirubin (***); *p* = 3.76 × 10⁻^4^ for 50 nM indirubin (***); *p* = 4.54 × 10⁻^4^ for 100 nM indirubin (***). **i**, **j**, **k**, **l**, Calculation of EC₅₀ for FICZ (**i**, **k**) and indirubin (**j**, **l**) determined by nonlinear regression of fold change values from the qRT-PCR experiments shown in **e-h** using log-transformed concentrations. **i**, EC_50_ = 2431 pM. **j**, EC_50_ = 392.6 pM. **k**, EC_50_ = 548.2 pM. **l**, EC_50_ = 135.0 pM.

**Extended Data Fig. 4.**
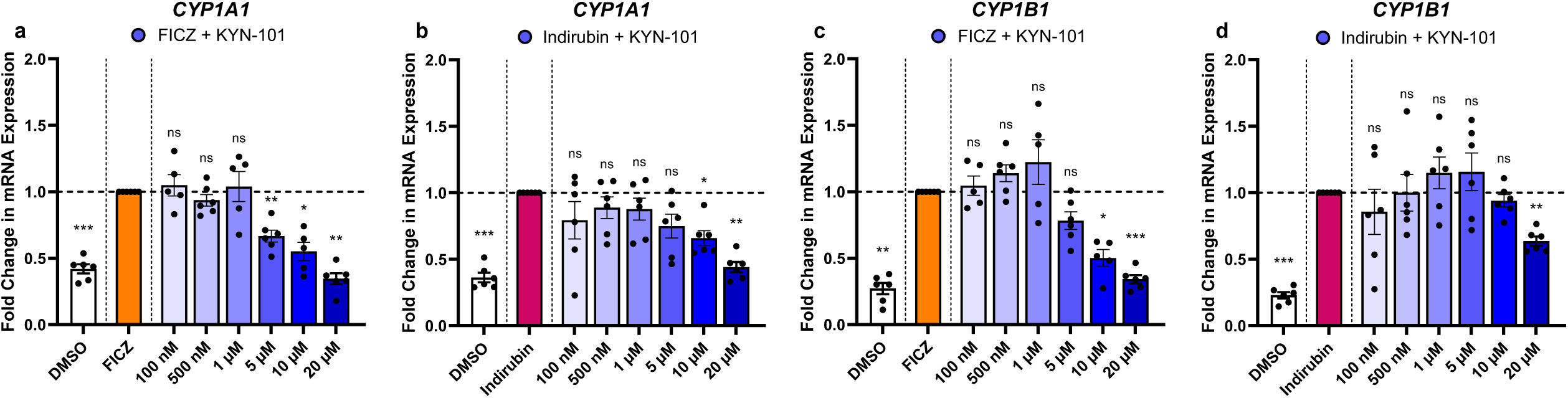
The AHR antagonist KYN-101 inhibits ligand-induced expression of AHR target genes. **a,** Quantification of *CYP1A1* mRNA expression in HEK293T cells following FICZ and KYN-101 treatment, measured by qRT-PCR and expressed as fold change relative to FICZ alone (set to 1). RNA was extracted following BiFC imaging, corresponding to Fig. 3d. *p* = 9.47 × 10 ^4^ for DMSO (***); *p* = 0.88 for 100 nM KYN-101 (ns); *p* = 0.45 for 500 nM KYN-101 (ns); *p* = 0.93 for 1 µM KYN-101 (ns); *p* = 8.21 × 10 ^3^ for 5 µM KYN-101 (**); *p* = 3.45 × 10 ^2^ for 10 µM KYN-101 (*); *p* = 2.80 × 10 ^3^ for 20 µM KYN-101 (**). **b,** Quantification of *CYP1A1* mRNA expression in HEK293T cells following indirubin and KYN-101 treatment, measured by qRT-PCR and expressed as fold change relative to indirubin alone (set to 1). RNA was extracted following BiFC imaging, corresponding to Fig. 3i. *p* = 6.94 × 10 ^4^ for DMSO (***); *p* = 0.45 for 100 nM KYN-101 (ns); *p* = 0.45 for 500 nM KYN-101 (ns); *p* = 0.45 for 1 µM KYN-101 (ns); *p* = 0.19 for 5 µM KYN-101 (ns); *p* = 1.30 × 10 ^2^ for 10 µM KYN-101 (*); *p* = 1.20 × 10 ^3^ for 20 µM KYN-101 (**). **c,** Quantification of *CYP1B1* mRNA expression in HEK293T cells following FICZ and KYN-101 treatment, measured by qRT-PCR and expressed as fold change relative to FICZ alone (set to 1). RNA was extracted following BiFC imaging, corresponding to Fig. 3d. *p* = 4.83 × 10 ^3^ for DMSO (**); *p* = 0.64 for 100 nM KYN-101 (ns); *p* = 0.23 for 500 nM KYN-101 (ns); *p* = 0.56 for 1 µM KYN-101 (ns); *p* = 0.11 for 5 µM KYN-101 (ns); *p* = 4.11 × 10 ^2^ for 10 µM KYN-101 (*); *p* = 4.46 × 10 ^4^ for 20 µM KYN-101 (***). **d,** Quantification of *CYP1B1* mRNA expression in HEK293T cells following indirubin and KYN-101 treatment, measured by qRT-PCR and expressed as fold change relative to indirubin alone (set to 1). RNA was extracted following BiFC imaging, corresponding to Fig. 3i. *p* = 3.08 × 10 ^4^ for DMSO (***); *p* = 0.76 for 100 nM KYN-101 (ns); *p* = 0.76 for 500 nM KYN-101 (ns); *p* = 0.76 for 1 µM KYN-101 (ns); *p* = 0.76 for 5 µM KYN-101 (ns); *p* = 0.72 for 10 µM KYN-101 (ns); *p* = 1.92 × 10 ^3^ for 20 µM KYN-101 (**). Data are mean ± SEM of n ≥ 5 (**a**, **c**) or n = 6 (**b**, **d**) independent biological replicates. Statistical significance was assessed using one-sample *t*-test on log-transformed fold-change values, followed by Holm-Šidák correction for multiple comparisons (**a-d**). KYN-101 treatment was performed in presence of 100 nM FICZ (**a**, **c**) or 100 nM indirubin (**b**, **d**).

**Extended Data Fig. 5.**
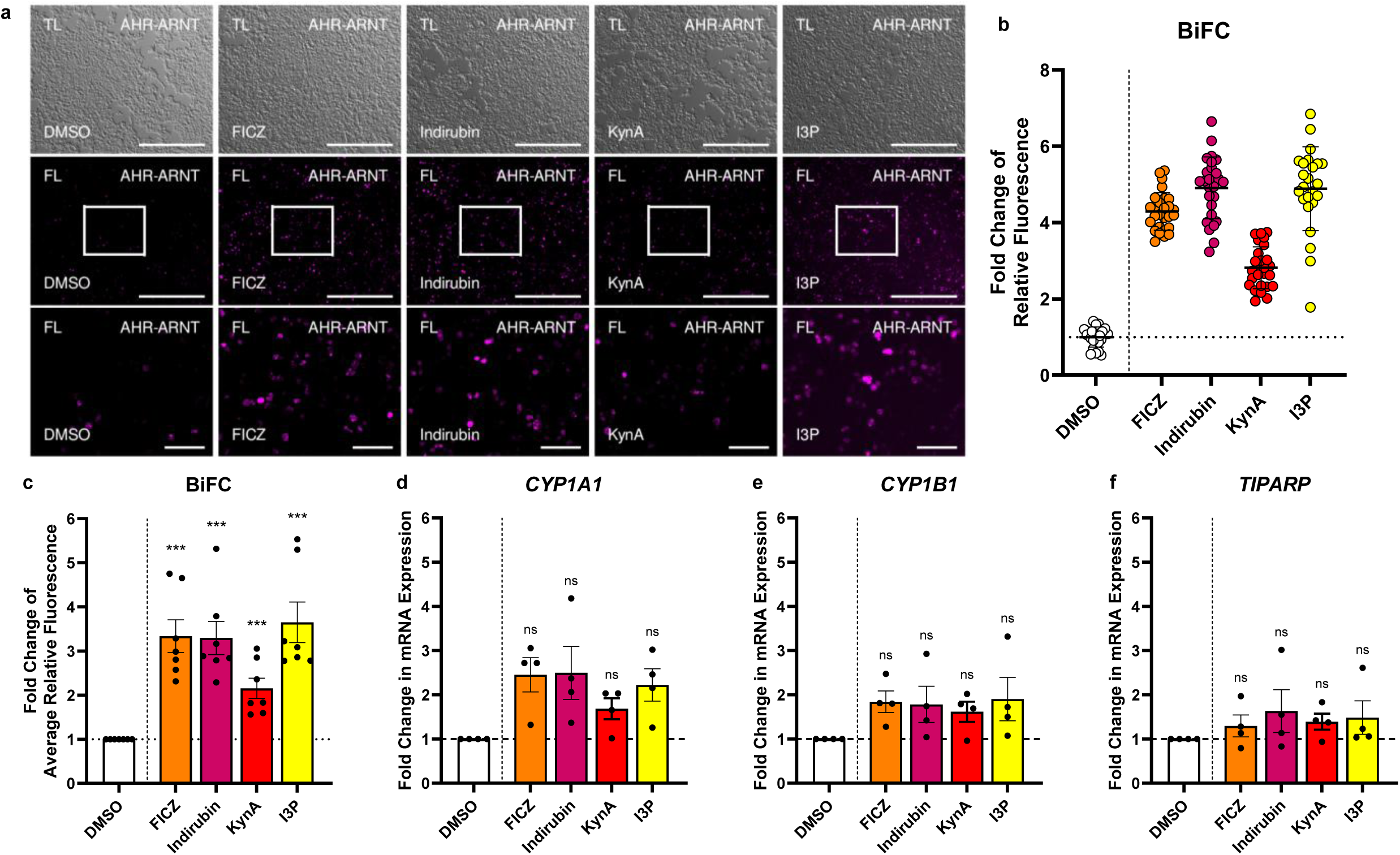
Additional AHR agonists induce AHR-ARNT interaction in HEK293T cells. **a,** Representative BiFC microscopy images in HEK293T cells. Top, transmitted light (TL); middle, fluorescence (FL); bottom, magnified views of the boxed regions in the FL images. Fluorescence signals are pseudocolored in magenta for visualization. Scale bars, 500 µm (top and middle panels) and 100 µm (bottom panel). **b,** Representative BiFC experiment in HEK293T cells showing normalized fluorescence-positive area upon treatment with FICZ, indirubin, KynA or I3P expressed as fold change relative to DMSO (average = 1). Each dot represents a single image (with an independent, non-overlapping field of view). Data are shown as mean ± SD of n ≥ 23 images per condition. **c,** Quantification of BiFC fluorescence (example shown in **b**). The average fluorescence per condition was calculated and normalized to DMSO (set to 1). *p* = 1.17 × 10 ^4^ for FICZ (***); *p* = 1.17 × 10 ^4^ for indirubin (***); *p* = 3.93 × 10 ^4^ for KynA (***); *p* = 1.17 × 10 ^4^ for I3P (***). **d, e, f,** Quantification of *CYP1A1* (**d**), *CYP1B1* (**e**) and *TIPARP* (**f**) mRNA expression in non-transfected HEK293T cells following agonist treatment with FICZ, indirubin, KynA or I3P, measured by qRT-PCR and expressed as fold change relative to DMSO (set to 1). **d**, *p* = 8.39 × ⁻ for FICZ (ns); *p* = 8.39 × 10⁻ for indirubin (ns); *p* = 8.39 × 10⁻ for KynA (ns); *p* = 8.39 × ⁻ for I3P (ns). **e**, *p* = 8.81 × 10⁻ for FICZ (ns); *p* = 0.21 for indirubin (ns); *p* = 0.21 for KynA (ns); *p* = 0.21 for I3P (ns). **f**, *p* = 0.57 for FICZ (ns); *p* = 0.57 for indirubin (ns); *p* = 0.39 for KynA (ns); *p* = 0.57 for I3P (ns). Treatment has been conducted with 100 nM FICZ or indirubin, 100 µM KynA or 25 µM I3P (**a-f**). Data are mean ± SEM of n = 7 (**c**) or n = 4 (**d-f**) independent biological replicates. Statistical significance was assessed using one-sample *t*-test on log-transformed fold-change values, followed by Holm-Šidák correction for multiple comparisons (**c-f**).

## References

1 Rothhammer, V. & Quintana, F. J. The aryl hydrocarbon receptor: an environmental sensor integrating immune responses in health and disease. Nature Reviews Immunology 19, 184–197 (2019). 10.1038/s41577-019-0125-8

2 Coumoul, X. et al. The aryl hydrocarbon receptor: structure, signaling, physiology and pathology. Signal Transduct Target Ther 11, 20 (2026). 10.1038/s41392-025-02500-8

3 Gutiérrez-Vázquez, C. & Quintana, F. J. Regulation of the Immune Response by the Aryl Hydrocarbon Receptor. Immunity 48, 19–33 (2018). 10.1016/j.immuni.2017.12.012

4 Lamas, B., Natividad, J. M. & Sokol, H. Aryl hydrocarbon receptor and intestinal immunity. Mucosal Immunology 11, 1024–1038 (2018). 10.1038/s41385-018-0019-2

5 Stockinger, B., Diaz, O. E. & Wincent, E. The influence of AHR on immune and tissue biology. EMBO Mol Med 16, 2290–2298 (2024). 10.1038/s44321-024-00135-w

6 Burbach, K. M., Poland, A. & Bradfield, C. A. Cloning of the Ah-receptor cDNA reveals a distinctive ligand-activated transcription factor. Proc Natl Acad Sci U S A 89, 8185–8189 (1992). 10.1073/pnas.89.17.8185

7 Fukunaga, B. N., Probst, M. R., Reisz-Porszasz, S. & Hankinson, O. Identification of functional domains of the aryl hydrocarbon receptor. J Biol Chem 270, 29270–29278 (1995). 10.1074/jbc.270.49.29270

8 Tumova, S., Dolezel, D. & Jindra, M. Conserved and Unique Roles of bHLH-PAS Transcription Factors in Insects - From Clock to Hormone Reception. J Mol Biol, 168332 (2023). 10.1016/j.jmb.2023.168332

9 Coumailleau, P., Poellinger, L., Gustafsson, J. A. & Whitelaw, M. L. Definition of a minimal domain of the dioxin receptor that is associated with Hsp90 and maintains wild type ligand binding affinity and specificity. J Biol Chem 270, 25291–25300 (1995). 10.1074/jbc.270.42.25291

10 Perdew, G. H. Association of the Ah receptor with the 90-kDa heat shock protein. J Biol Chem 263, 13802–13805 (1988).

11 Meyer, B. K., Pray-Grant, M. G., Vanden Heuvel, J. P. & Perdew, G. H. Hepatitis B virus X-associated protein 2 is a subunit of the unliganded aryl hydrocarbon receptor core complex and exhibits transcriptional enhancer activity. Mol Cell Biol 18, 978–988 (1998). 10.1128/mcb.18.2.978

12 Kazlauskas, A., Poellinger, L. & Pongratz, I. Evidence that the co-chaperone p23 regulates ligand responsiveness of the dioxin (Aryl hydrocarbon) receptor. J Biol Chem 274, 13519–13524 (1999). 10.1074/jbc.274.19.13519

13 Hankinson, O. The aryl hydrocarbon receptor complex. Annu Rev Pharmacol Toxicol 35, 307–340 (1995). 10.1146/annurev.pa.35.040195.001515

14 Jain, S., Dolwick, K. M., Schmidt, J. V. & Bradfield, C. A. Potent transactivation domains of the Ah receptor and the Ah receptor nuclear translocator map to their carboxyl termini. J Biol Chem 269, 31518–31524 (1994).

15 Wu, D., Potluri, N., Kim, Y. & Rastinejad, F. Structure and dimerization properties of the aryl hydrocarbon receptor PAS-A domain. Mol Cell Biol 33, 4346–4356 (2013). 10.1128/mcb.00698-13

16 Larigot, L., Juricek, L., Dairou, J. & Coumoul, X. AhR signaling pathways and regulatory functions. Biochim Open 7, 1–9 (2018). 10.1016/j.biopen.2018.05.001

17 Sadik, A. et al. IL4I1 Is a Metabolic Immune Checkpoint that Activates the AHR and Promotes Tumor Progression. Cell 182, 1252–1270 e1234 (2020). 10.1016/j.cell.2020.07.038

18 Poland, A. P., Glover, E., Robinson, J. R. & Nebert, D. W. Genetic expression of aryl hydrocarbon hydroxylase activity. Induction of monooxygenase activities and cytochrome P1-450 formation by 2,3,7,8-tetrachlorodibenzo-p-dioxin in mice genetically “nonresponsive” to other aromatic hydrocarbons. J Biol Chem 249, 5599–5606 (1974).

19 Sutter, T. R. et al. Complete cDNA sequence of a human dioxin-inducible mRNA identifies a new gene subfamily of cytochrome P450 that maps to chromosome 2. J Biol Chem 269, 13092–13099 (1994).

20 Pollenz, R. S. The mechanism of AH receptor protein down-regulation (degradation) and its impact on AH receptor-mediated gene regulation. Chem Biol Interact 141, 41–61 (2002). 10.1016/s0009-2797(02)00065-0

21 Ma, Q. & Baldwin, K. T. 2,3,7,8-tetrachlorodibenzo-p-dioxin-induced degradation of aryl hydrocarbon receptor (AhR) by the ubiquitin-proteasome pathway. Role of the transcription activaton and DNA binding of AhR. J Biol Chem 275, 8432–8438 (2000). 10.1074/jbc.275.12.8432

22 Dixon, A. S. et al. NanoLuc Complementation Reporter Optimized for Accurate Measurement of Protein Interactions in Cells. ACS Chem Biol 11, 400–408 (2016). 10.1021/acschembio.5b00753

23 Hu, C. D., Chinenov, Y. & Kerppola, T. K. Visualization of interactions among bZIP and Rel family proteins in living cells using bimolecular fluorescence complementation. Mol Cell 9, 789–798 (2002). 10.1016/s1097-2765(02)00496-3

24 Rannug, U. et al. Structure elucidation of two tryptophan-derived, high affinity Ah receptor ligands. Chem Biol 2, 841–845 (1995). 10.1016/1074-5521(95)90090-x

25 Adachi, J. et al. Indirubin and indigo are potent aryl hydrocarbon receptor ligands present in human urine. J Biol Chem 276, 31475–31478 (2001). 10.1074/jbc.C100238200

26 Powis, M., Celius, T. & Matthews, J. Differential ligand-dependent activation and a role for Y322 in aryl hydrocarbon receptor-mediated regulation of gene expression. Biochem Biophys Res Commun 410, 859–865 (2011). 10.1016/j.bbrc.2011.06.079

27 Oberg, M., Bergander, L., Håkansson, H., Rannug, U. & Rannug, A. Identification of the tryptophan photoproduct 6-formylindolo[3,2-b]carbazole, in cell culture medium, as a factor that controls the background aryl hydrocarbon receptor activity. Toxicol Sci 85, 935–943 (2005). 10.1093/toxsci/kfi154

28 Veldhoen, M., Hirota, K., Christensen, J., O’Garra, A. & Stockinger, B. Natural agonists for aryl hydrocarbon receptor in culture medium are essential for optimal differentiation of Th17 T cells. J Exp Med 206, 43–49 (2009). 10.1084/jem.20081438

29 Pollenz, R. S., Sattler, C. A. & Poland, A. The aryl hydrocarbon receptor and aryl hydrocarbon receptor nuclear translocator protein show distinct subcellular localizations in Hepa 1c1c7 cells by immunofluorescence microscopy. Molecular Pharmacology 45, 428 (1994).

30 Campesato, L. F. et al. Blockade of the AHR restricts a Treg-macrophage suppressive axis induced by L-Kynurenine. Nat Commun 11, 4011 (2020). 10.1038/s41467-020-17750-z

31 Schwarcz, R., Bruno, J. P., Muchowski, P. J. & Wu, H. Q. Kynurenines in the mammalian brain: when physiology meets pathology. Nat Rev Neurosci 13, 465–477 (2012). 10.1038/nrn3257

32 Bittinger, M. A., Nguyen, L. P. & Bradfield, C. A. Aspartate aminotransferase generates proagonists of the aryl hydrocarbon receptor. Mol Pharmacol 64, 550–556 (2003). 10.1124/mol.64.3.550

33 Diani-Moore, S. et al. Identification of the aryl hydrocarbon receptor target gene TiPARP as a mediator of suppression of hepatic gluconeogenesis by 2,3,7,8-tetrachlorodibenzo-p-dioxin and of nicotinamide as a corrective agent for this effect. J Biol Chem 285, 38801–38810 (2010). 10.1074/jbc.M110.131573

34 Matthews, J. AHR toxicity and signaling: Role of TIPARP and ADP-ribosylation. Current Opinion in Toxicology 2, 50–57 (2017). 10.1016/j.cotox.2017.01.013

35 Kolluri, S. K., Jin, U. H. & Safe, S. Role of the aryl hydrocarbon receptor in carcinogenesis and potential as an anti-cancer drug target. Arch Toxicol 91, 2497–2513 (2017). 10.1007/s00204-017-1981-2

36 Gradin, K. et al. Functional interference between hypoxia and dioxin signal transduction pathways: competition for recruitment of the Arnt transcription factor. Mol Cell Biol 16, 5221–5231 (1996). 10.1128/mcb.16.10.5221

37 Vorrink, S. U. & Domann, F. E. Regulatory crosstalk and interference between the xenobiotic and hypoxia sensing pathways at the AhR-ARNT-HIF1α signaling node. Chem Biol Interact 218, 82–88 (2014). 10.1016/j.cbi.2014.05.001

38 Shafighi, M. et al. Dimethyloxalylglycine stabilizes HIF-1α in cultured human endothelial cells and increases random-pattern skin flap survival in vivo. Plast Reconstr Surg 128, 415–422 (2011). 10.1097/PRS.0b013e31821e6e69

39 Gruszczyk, J. et al. Cryo-EM structure of the agonist-bound Hsp90-XAP2-AHR cytosolic complex. Nature Communications 13, 7010 (2022). 10.1038/s41467-022-34773-w

40 Diao, X. et al. Structural basis for the ligand-dependent activation of heterodimeric AHR-ARNT complex. Nat Commun 16, 1282 (2025). 10.1038/s41467-025-56574-7

41 Lamas, B. et al. CARD9 impacts colitis by altering gut microbiota metabolism of tryptophan into aryl hydrocarbon receptor ligands. Nat Med 22, 598–605 (2016). 10.1038/nm.4102

42 Stockinger, B., Shah, K. & Wincent, E. AHR in the intestinal microenvironment: safeguarding barrier function. Nat Rev Gastroenterol Hepatol 18, 559–570 (2021). 10.1038/s41575-021-00430-8

43 Roager, H. M. & Licht, T. R. Microbial tryptophan catabolites in health and disease. Nat Commun 9, 3294 (2018). 10.1038/s41467-018-05470-4

44 Takamura, T. et al. Activation of the aryl hydrocarbon receptor pathway may ameliorate dextran sodium sulfate-induced colitis in mice. Immunol Cell Biol 88, 685–689 (2010). 10.1038/icb.2010.35

45 Rannug, A. How the AHR Became Important in Intestinal Homeostasis-A Diurnal FICZ/AHR/CYP1A1 Feedback Controls Both Immunity and Immunopathology. Int J Mol Sci 21 (2020). 10.3390/ijms21165681

46 Sugihara, K. et al. Aryl hydrocarbon receptor-mediated induction of microsomal drug-metabolizing enzyme activity by indirubin and indigo. Biochem Biophys Res Commun 318, 571–578 (2004). 10.1016/j.bbrc.2004.04.066

47 Peter Guengerich, F., et al. Aryl hydrocarbon receptor response to indigoids in vitro and in vivo. Arch Biochem Biophys 423, 309–316 (2004). 10.1016/j.abb.2004.01.002

48 Spink, D. C. et al. Estrogen regulates Ah responsiveness in MCF-7 breast cancer cells. Carcinogenesis 24, 1941–1950 (2003). 10.1093/carcin/bgg162

49 Farmahin, R., Crump, D., O’Brien, J. M., Jones, S. P. & Kennedy, S. W. Time-dependent transcriptomic and biochemical responses of 6-formylindolo[3,2-b]carbazole (FICZ) and 2,3,7,8-tetrachlorodibenzo-p-dioxin (TCDD) are explained by AHR activation time. Biochem Pharmacol 115, 134–143 (2016). 10.1016/j.bcp.2016.06.005

50 Rannug, U., Bramstedt, H. & Nilsson, U. The presence of genotoxic and bioactive components in indigo dyed fabrics--a possible health risk? Mutat Res 282, 219–225 (1992). 10.1016/0165-7992(92)90099-4

51 Karabogdan, I. et al. Exploration of Agonist and Antagonist Binding Sites within the Cytosolic AHR Complex Using Molecular Modeling. ACS Omega 11, 16070–16087 (2026). 10.1021/acsomega.5c10598

52 Opitz, C. A., Holfelder, P., Prentzell, M. T. & Trump, S. The complex biology of aryl hydrocarbon receptor activation in cancer and beyond. Biochem Pharmacol 216, 115798 (2023). 10.1016/j.bcp.2023.115798

53 Wright, E. J., De Castro, K. P., Joshi, A. D. & Elferink, C. J. Canonical and non-canonical aryl hydrocarbon receptor signaling pathways. Curr Opin Toxicol 2, 87–92 (2017). 10.1016/j.cotox.2017.01.001

54 Sakurai, S., Shimizu, T. & Ohto, U. The crystal structure of the AhRR–ARNT heterodimer reveals the structural basis of the repression of AhR-mediated transcription. Journal of Biological Chemistry 292, 17609–17616 (2017). 10.1074/jbc.M117.812974

55 Janssens, L. K. & Stove, C. P. Sensing an Oxygen Sensor: Development and Application of Activity-Based Assays Directly Monitoring HIF Heterodimerization. Anal Chem 93, 14462–14470 (2021). 10.1021/acs.analchem.1c02923

56 Janssens, L. K. & Stove, C. P. The ‘ABC’ of split-nanoluciferase HIF heterodimerization bioassays: Applications, Benefits & Considerations. Biochem Pharmacol 229, 116478 (2024). 10.1016/j.bcp.2024.116478

57 Weiler, M. et al. mTOR target NDRG1 confers MGMT-dependent resistance to alkylating chemotherapy. Proc Natl Acad Sci U S A 111, 409–414 (2014). 10.1073/pnas.1314469111

58 Prentzell, M. T. et al. G3BPs tether the TSC complex to lysosomes and suppress mTORC1 signaling. Cell 184, 655–674.e627 (2021). 10.1016/j.cell.2020.12.024

59 Tumova, S. & Jindra, M. Ligand-dependent protein interactions of the juvenile hormone receptor captured in real time. Febs j 290, 2881–2894 (2023). 10.1111/febs.16719

60 Chu, J. et al. A novel far-red bimolecular fluorescence complementation system that allows for efficient visualization of protein interactions under physiological conditions. Biosens Bioelectron 25, 234–239 (2009). 10.1016/j.bios.2009.06.008

61 Schmittgen, T. D. & Livak, K. J. Analyzing real-time PCR data by the comparative C(T) method. Nat Protoc 3, 1101–1108 (2008). 10.1038/nprot.2008.73

