## Supplementary Data 1 for "Capturing early events in aryl hydrocarbon receptor activation using two complementary protein-protein interaction assays"

**Supplementary Data 1 – Reagents and Tools Table**

| **Reagent/Resource** | | **Reference or Source** | | | **Identifier or Catalog Number** | |
| --- | --- | --- | --- | --- | --- | --- |
| **Experimental Models** | | | | | | |
| CHO-K1 (Cricetulus griseus) | | American Type Cell Culture Collection (ATCC) | | | CCL-61 | |
| HEK293T (*H. sapiens*) | | American Type Cell Culture Collection (ATCC) | | | CRL-3216 | |
| LN-229 (*H. sapiens*) | | American Type Cell Culture Collection (ATCC) | | | CRL-2611 | |
| **Antibodies** | | | | | | |
| Anti-HA High Affinity | | Roche | | | 11867423001 | |
| c-Myc (9E10) | | Thermo Fisher Scientific | | | MA1-980 | |
| DYKDDDDK (FLAG) Tag [FG4R] | | Thermo Fisher Scientific | | | MA1-91878 | |
| Goat anti-mouse IgG Secondary Antibody, HRP-coupled | | Thermo Fisher Scientific | | | 31430 | |
| Goat anti-rabbit IgG Secondary Antibody, HRP-coupled | | Thermo Fisher Scientific | | | 31460 | |
| Goat anti-rat IgG  Secondary Antibody, HRP-coupled | | Thermo Fisher Scientific | | | 31470 | |
| Myc-Tag [9B11] | | Cell Signaling Technology | | | 2276 | |
| Peroxidase AffiniPure® F(ab')₂ Fragment Goat Anti-Mouse IgG | | Jackson ImmunoResearch Laboratories, Inc | | | 115-036-003 | |
| Peroxidase AffiniPure® F(ab')₂ Fragment Goat Anti-Rabbit IgG | | Jackson ImmunoResearch Laboratories, Inc | | | 111-036-003 | |
| S6 Ribosomal Protein (S6RP) [5G10] | | Cell Signaling Technology | | | 2217 | |
| TUBA1B (Tubulin) [EPR1333] | | Abcam | | | ab108629 | |
| **Protein Sequences** | | | | | | |
| **Name** | **Uniprot-ID** | | **Gene-ID** | **CCDS-ID** | | **Length** |
| AHR | P35869 | | 196 | 5366.1 | | M1-L848 |
| ARNT | P27540 | | 405 | 970.1 | | M1-E789 |
| HSP90 (HSP90AB1) | P08238 | | 3326 | 4909.1 | | M1-D724 |
| **Vectors and Plasmids** | | | | | | |
| AHR Human Tagged ORF Clone (NM_001621) | | OriGene Technologies | | | RG209832 | |
| ARNT Human Tagged ORF Clone (NM_001668.4) | | OriGene Technologies | | | RG216724 | |
| HSP90AB1 - HaloTag® human ORF in pFN21A | | Promega | | | FHC01815 | |
| pBiT1.1-N [TK/LgBiT] | | Promega | | | N198A | |
| pBiT2.1-N [TK/SmBiT] | | Promega | | | N199A | |
| pBiT1.1-N-Lg-myc-AHR | | This paper | | |  | |
| pBiT2.1-N-Sm-FLAG-ARNT | | This paper | | |  | |
| pBiT2.1-N-Sm-myc-AHR-WT | | This paper | | |  | |
| pBiT2.1-N-Sm-myc-AHR-Y322 | | This paper | | |  | |
| pBiT1.1-N-Lg-FLAG-ARNT | | This paper | | |  | |
| pBiT1.1-N-Lg-FLAG-HSP90 | | This paper | | |  | |
| pGL4.43 [luc2P/XRE/Hygro] Vector | | Promega | | | E4121 | |
| pGL4.74 [hRluc/TK] Vector | | Promega | | | E6921 | |
| pGW-MYC-LC151 | | Stefan Pusch^1^ | | |  | |
| pGW-HA-LN151 | | Stefan Pusch^1^ | | |  | |
| pDONR201 | | Clone repository of the DKFZ Genomics and Proteomics Core Facility (GPCF) | | |  | |
| pDONR201-AHR | | This paper | | |  | |
| pDONR201-ARNT | | This paper | | |  | |
| pDONR223-HSP90 | | Clone repository of the DKFZ Genomics and Proteomics Core Facility (GPCF) | | |  | |
| pGW-AHR-HA-LN151 | | This paper | | |  | |
| pGW-ARNT-MYC-LC151 | | This paper | | |  | |
| pGW-HSP90-MYC-LC151 | | This paper | | |  | |
| **Bacterial strains** | | | | | | |
| DB3.1 (*Escherichia coli*) | | Thermo Fisher Scientific | | | 11782018 (discontinued) | |
| DH5α (*Escherichia coli*) | | New England Biolabs | | | C2987H | |
| Stbl3 (*Escherichia coli*) | | Thermo Fisher Scientific | | | C737303 | |
| **Chemicals and Kits** | | | | | | |
| (2Z)-Indirubin | | Sigma Aldrich | | | SML0280 | |
| 4,6-Diamino-2-phenylindol-2HCl (DAPI) | | SERVA Electrophoresis GmbH | | | 18860 | |
| 6-Formylindolo[3,2-b]carbazole (FICZ) | | MedChemExpress LLC | | | HY-12451 | |
| 6-Formylindolo[3,2-b]carbazole (FICZ) | | Sigma Aldrich | | | SML1489 | |
| β-Mercaptoethanol | | Sigma Aldrich | | | M3148 | |
| β-Mercaptoethanol | | Serva | | | 28625 | |
| Albumin Bovine, Fraction V | | Serva | | | 11926 | |
| Amino Extract Buffer (IDK® Amino Extract) | | Immundiagnostik AG | | | K7999 | |
| Bovine Serum Albumin | | Carl Roth | | | 8076.3 | |
| Bromophenol Blue | | Sigma Aldrich | | | B5525 | |
| Complete Protease Inhibitor Cocktail | | Roche | | | 11836145001 | |
| Dimethyloxaloylglycine (DMOG) | | MedChemExpress LLC | | | HY-15893 | |
| Dimethylsulfoxid (DMSO) | | Sigma-Aldrich | | | D8418 | |
| Dual-Glo(R) Luciferase Assay System | | Promega | | | E2940 | |
| FuGENE® HD Transfection Reagent | | Promega | | | E2311 | |
| Glycerol | | Sigma-Aldrich | | | G5516 | |
| Glycine | | GERBU | | | 1023 | |
| HEPES | | VWR | | | 441485H | |
| High Capacity cDNA Reverse Transcription Kit | | Thermo Fisher Scientific | | | 4368814 | |
| Hydroxychloride Acid (HCl), 25% | | Carl Roth | | | X897.1 | |
| IGEPAL | | Sigma Aldrich | | | I8896 | |
| Indole-3-pyruvate (I3P) | | Sigma Aldrich | | | I7017 | |
| Isopropanol | | Thermo Fisher Scientific | | | P/7500/PC17 | |
| KYN-101 | | MedChemExpress LLC | | | HY-134217 | |
| Kynurenic Acid (KynA) | | Sigma Aldrich | | | K3375 | |
| LipofectamineTM 3000 Transfection Reagent-1.5 mL | | Thermo Fisher Scientific | | | L3000015 | |
| Methanol | | Thermo Fisher Scientific | | | M/4000/PC17 | |
| Nano-Glo® Live Cell Assay System | | Promega | | | N2012 | |
| Nonfat dried milk powder | | AppliChem | | | A0830 | |
| NucleoBond Xtra Midi | | Macherey-Nagel | | | 740410.50 | |
| OptiMEM | | Gibco | | | 11058-021 | |
| PageRuler™ Plus Prestained Protein Ladder, 10-250 kDa | | Thermo Fisher Scientific | | | 26620 | |
| Paraformaldehyde | | Sigma Aldrich | | | 16005 | |
| Passive Lysis 5X Buffer | | Promega | | | E194A | |
| Phosphatase Inhibitor Cocktail 2 | | Sigma Aldrich | | | P5726 | |
| Phosphatase Inhibitor Cocktail 3 | | Sigma Aldrich | | | P0044 | |
| Pierce™ ECL Western Blotting Substrate | | Thermo Fisher Scientific | | | 32106 | |
| Pierce™ LDS Sample Buffer, Non-Reducing (4X) | | Thermo Fisher Scientific | | | 84788 | |
| Precision Plus Protein™ Dual Color Standards | | Bio-Rad | | | 1610394 | |
| Protein Assay Dye Reagent Concentrate | | Bio-Rad | | | 5000006 | |
| RNase-Free DNase Set | | Qiagen | | | 79254 | |
| RNeasy Mini Kit | | Qiagen | | | 74106 | |
| Sodium Azide | | AppliChem | | | A1430,0100 | |
| Sodium Deoxycholate | | AppliChem | | | A1531 | |
| Sodium Dodecyl Sulfate (SDS) | | Carl Roth | | | 8029.3 | |
| Sodium Hydroxide (NaOH) | | Reagecon | | | 3006300 | |
| SuperSignal™ West Femto Maximum Sensitivity Substrate | | Thermo Fisher Scientific | | | 34096 | |
| SYBR Green PCR Master Mix | | Thermo Fisher Scientific | | | 4309155 | |
| Triton™ X-100 | | Sigma Aldrich | | | T8787 | |
| Trizma® base | | Sigma Aldrich | | | T1503 | |
| Tween-20 | | Sigma Aldrich | | | P9416 | |
| Tween-20 | | AppliChem | | | A1389 | |
| **Reagents for Cell Culture** | | | | | | |
| Dulbecco's Modified Eagle Medium (DMEM) w/ 4.5 g/L Glucose | | Gibco®, Thermo Fisher Scientific | | | 31053-028 | |
| Dulbecco's Modified Eagle Medium (DMEM) w/ 4.5 g/L Glucose, GlutaMAX™, pyruvate | | Gibco®, Thermo Fisher Scientific | | | 31966-021 | |
| Fetal Bovine Serum (FBS) | | Gibco®, Thermo Fisher Scientific | | | 26140079 | |
| Fetal Bovine Serum (FBS) (Heat inactivated) | | Sigma Aldrich | | | F4135 | |
| L-Glutamine | | Life Technologies GmbH | | | 25030-024 | |
| Mycoplasma Test Kit Venor®GeM Classic | | Minerva Biolabs GmbH | | | 11-1025 | |
| Penicillin-Streptomycin | | Thermo Fisher Scientific | | | 15140122 | |
| Phosphate-buffered Saline (PBS) | | Gibco®, Thermo Fisher Scientific | | | 14190094 | |
| Sodium Pyruvate | | Gibco®, Thermo Fisher Scientific | | | 11360 | |
| Trypan Blue Stain (0.4%) | | Invitrogen | | | T10282 | |
| Trypsin-EDTA (0.5%) | | Gibco®, Thermo Fisher Scientific | | | 15400054 | |
| **Consumables** | | | | | | |
| 0.5 mL Safe-Lock Tubes | | Eppendorf SE | | | 0030121023 | |
| 1.5 mL Safe-Lock Tubes | | Eppendorf SE | | | 0030120086 | |
| 2 mL Safe-Lock Tubes | | Eppendorf SE | | | 0030120094 | |
| 5 mL Serological Pipette | | Thermo Fisher Scientific | | | 15082 | |
| 5 mL Serological Pipette | | Jet Biofil | | | GSP010005 | |
| 6-well Cell Culture Plate | | Greiner Bio-One | | | 657160 | |
| 10 mL Serological Pipette | | Greiner Bio-One | | | 607180 | |
| 25 mL Serological Pipette | | Greiner Bio-One | | | 760180 | |
| 15 mL Conical Tube | | Greiner Bio-One | | | 188271-N | |
| 50 mL Conical Tube | | Greiner Bio-One | | | 227261 | |
| 96-well PCR Platte, Halbrand, 0.2 ml/well | | Steinbrenner Laborsysteme GmbH | | | SL-PP96-2 | |
| µ-Plate 24 Well | | ibidi GmbH | | | 82426 | |
| Amersham™ Protran Premium 0.45 NC Nitrocellulose Western Blotting Membranes | | Cytiva | | | 10600003 | |
| Corning® 96-well Solid White Flat Bottom Polystyrene TC-treated Microplates | | Corning Life Sciences | | | 3917 | |
| Countess™ Cell Counting Chamber Slides | | Invitrogen | | | C10312 | |
| MicroAmpTM Optical Adhesive Film-100 Covers | | Thermo Fisher Scientific | | | 4311971 | |
| Neptune 10 µL Extended Length Universal Barrier Tip | | Neptune Scientific | | | BT10XLS3 | |
| Neptune 200 µL Universal Barrier Tip | | Neptune Scientific | | | BT200 | |
| Neptune 1000 µL Universal Barrier Tip | | Neptune Scientific | | | BT1000.96 | |
| Nunc F96 MicroWell Delta Surface (white) | | Thermo Fisher Scientific | | | 10072151 | |
| PCR 8-Tube Strips | | Greiner Bio-One | | | 671221 | |
| Polyvinylidene Difluoride (PVDF) Membranes for Immunoblotting (Pore Size 0.45 µM) | | Merck Millipore | | | IPVH00010 | |
| Sartorius® MinisartNML 0.2µm, 28mm FX eo-ster, 50pc | | Sartorius | | | S7597------FXOSK | |
| T175 Cell Culture Flask | | Greiner Bio-One | | | 159900 | |
| T75 Cell Culture Flask | | Greiner Bio-One | | | 658170 | |
| T25 Nunc EasYFlask Cell Culture Flask | | Thermo Fisher Scientific | | | 156367 | |
| Thermo Scientific™ Nunc™ MicroWell™ 96-Well, Nunclon Delta-Treated, Flat-Bottom Microplate | | Thermo Fisher Scientific | | | 136101 | |
| TipOne® 10/20 µL XL Graduated Filter Tip | | Starlab | | | S1120-3710 | |
| TipOne® 1000 µL XL Graduated Filter Tip | | Starlab | | | S1122-1730 | |
| TipOne® 20 µL  Bevelled Filter Tip | | Starlab | | | S1120-1710 | |
| TipOne® 200 µL  Graduated Filter Tip | | Starlab | | | S1120-8710 | |
| TipOne® 200 µL Yellow Tip | | Starlab | | | S1111-0706 | |
| Type F Immersion Liquid | | Thermo Fisher Scientific | | | 11944399 | |
| Whatman Grade 3MM Chr Blotting Paper | | Cytiva | | | 3030-917 | |
| **Instruments** | | | | | | |
| Alliance Q9 Luminescent Image Analyzer | | UVItec Cambridge Bio Imaging SAS | | | DEV-ALLI-Q9 | |
| Biometra T3000 thermocycler | | Analytik Jena GmbH + Co. KG | | |  | |
| Cell Culture Incubator | | SANYO Electric Co., Ltd. | | | MCO-20AIC CO2 Incubator | |
| Dual Stacked Water Jacketed CO_2_ Incubator | | Shel Lab | | | Model 2424/2 | |
| ChemiDoc MP Imaging System | | Bio-Rad | | | 12003154 | |
| ChemiDoc XRS+ Camera System | | Bio-Rad | | | 1708265 | |
| CLARIOstar® | | BMG Labtech | | |  | |
| Countess 3 Cell Counter | | Thermo Fisher Scientific | | |  | |
| FastGene NanoSpec Photometer | | NIPPON Genetics EUROPE GmbH | | |  | |
| Maxisafe 2030i Biological Safety Cabinet (Class II) | | Thermo Fisher Scientific | | |  | |
| MICA Microhub | | Leica Microsystems | | |  | |
| Orion II Microplate Luminometer | | Berthold Technologies | | | Orion II LB 965 | |
| QuantStudio 3 Real-Time PCR System | | Thermo Fisher Scientific | | |  | |
| Telstar BIO II A Class II (Type A2) Biological Safety Cabinet | | Telstar | | |  | |
| Ultrasonic bath “Digital 10 P” | | Sonorex | | |  | |
| Ultrospec 2100 pro spectrophotometer | | Amersham Biosciences Corp. | | |  | |
| **Software** | | | | | | |
| CLARIOstar® Reader Control Software | | BMG Labtech  https://www.bmglabtech.com/de/microplate-reader-software/ | | | V5.40 R2 | |
| Design & Analysis Software | | Thermo Fisher Scientific  https://www.thermofisher.com/de/de/home/technical-resources/software-downloads/quantstudio-3-5-real-time-pcr-systems.html | | | 2.7.0 | |
| GraphPad Prism | | Dotmatics  https://www.graphpad.com/scientific-software/prism/ | | | v.10.6.1 | |
| ImageJ | | National Institute of Health  https://imagej.net/ij/ | | | 1.54f | |
| Image Lab Software | | Bio-Rad  https://www.bio-rad.com/de-de/product/image-lab-software?ID=KRE6P5E8Z | | | v.6.0.1 | |
| LAS X Office | | Leica Microsystems CMS GmbH  https://www.leica-microsystems.com/de/produkte/mikroskop-software/p/leica-las-x-ls/downloads/ | | | 1.4.7.28982 | |
| MARS Data Analysis Software | | BMG Labtech  https://www.bmglabtech.com/de/microplate-reader-software/ | | | 3.31 | |
| Microsoft Excel | | Microsoft  https://www.microsoft.com/de-de | | | 2016 | |
| Microsoft Power Point | | Microsoft  https://www.microsoft.com/de-de | | | 2016 | |
| Microsoft Word | | Microsoft  https://www.microsoft.com/de-de | | | 2016 | |
| Open Microscopy Environment Remote Objects (OMERO).insight | | The Open Microscopy Environment  https://www.openmicroscopy.org/omero/downloads/ | | | 5.8.3 | |
| Open Microscopy Environment Remote Objects (OMERO).figure | | The Open Microscopy Environment  https://www.openmicroscopy.org/omero/figure/ | | | 7.2.1 | |
| SnapGene | | Dotmatics  https://www.snapgene.com/ | | | v8.2.0 | |

| **Primer** | | |
| --- | --- | --- |
| **qRT-PCR** | | |
| *18S RNA* | GATGGGCGGCGGAAAATAG | GCGTGGATTCTGCATAATGGT |
| *CYP1A1* | CCCCCACAGCACAACAAGAG | GGGTGAGAAACCGTTCAGGT |
| *CYP1B1* | GACGCCTTTATCCTCTCTGCG | ACGACCTGATCCAATTCTGCC |
| *TIPARP* | CACCCTCTAGCAATGTCAAC | CAGACTCGGGATACTCTCTC |
| **Gateway cloning** | | |
| AHR_AttB1 | CAAAAAAGCAGGCTCCACCATGAACAGCAGCAGCGCCAAC | |
| AHR_AttB2_m | CAAGAAAGCTGGGTTTTACAGGAATCCACTGGATGTCAAATC | |
| AHR_AttB2_o | CAAGAAAGCTGGGTTCAGGAATCCACTGGATGTCAAATCAG | |
| ARNT_AttB1 | CAAAAAAGCAGGCTCCACCATGGCGGCGACTACTGCCAAC | |
| ARNT_AttB2_m | CAAGAAAGCTGGGTTCTATTCTGAAAAGGGGGGAAACATAGTTAG | |
| ARNT_AttB2_o | CAAGAAAGCTGGGTTTTCTGAAAAGGGGGGAAACATAGTTAGATC | |
| AttB1 | GGGGACAAGTTTGTACAAAAAAGCAGGCTCCACC | |
| AttB2 | GGGGACCACTTTGTACAAGAAAGCTGGGTT | |
| **Site-directed mutagenesis** | | |
| AHR Y322A | fwd: 5′- GCACGAGAGGCTCAGGT**GC**TCAGTTTATTCATGCAGCTG -3′  rev: 5′- CAGCTGCATGAATAAACTGA**GC**ACCTGAGCCTCTCGTGC -3′ | |
