## Supplementary Data 2 for "Capturing early events in aryl hydrocarbon receptor activation using two complementary protein-protein interaction assays"

### ImageJ macro for automated quantification of BiFC-mediated fluorescence

```
title=getTitle();
roiManager("Reset");
setSlice(1);
run("Duplicate...", "title=dupl channels=1");
run("Subtract Background...", "rolling=50");
run("Gaussian Blur...", "sigma=1");
setAutoThreshold("Default dark");
run("Threshold...");
setThreshold(150, 65535);
run("Set Measurements...", "integrated limit redirect=None decimal=3");
rename(title+" positive area");
run("Analyze Particles...", "size=20-Infinity summarize add");
close();
selectWindow(title);
setSlice(2);
run("Duplicate...", "title=dupl channels=2");
run("Variance...", "radius=10");
run("Enhance Contrast", "saturated=0.35");
run("Apply LUT");
setOption("ScaleConversions", true);
run("8-bit");
setThreshold(30, 255, "raw");
rename(title+" total area");
selectWindow("Threshold");
run("Analyze Particles...", "size=800-Infinity summarize");
run("Create Selection");
roiManager("Add");
close();
nRoi=roiManager("count");
roiManager("Show All without labels");
roiManager("Select", nRoi-1);
```
