## Supplementary Data 4 for "Capturing early events in aryl hydrocarbon receptor activation using two complementary protein-protein interaction assays"

**Vector maps of used NanoBiT and BiFC plasmids**

a) Vector map of NanoBiT AHR plasmid.

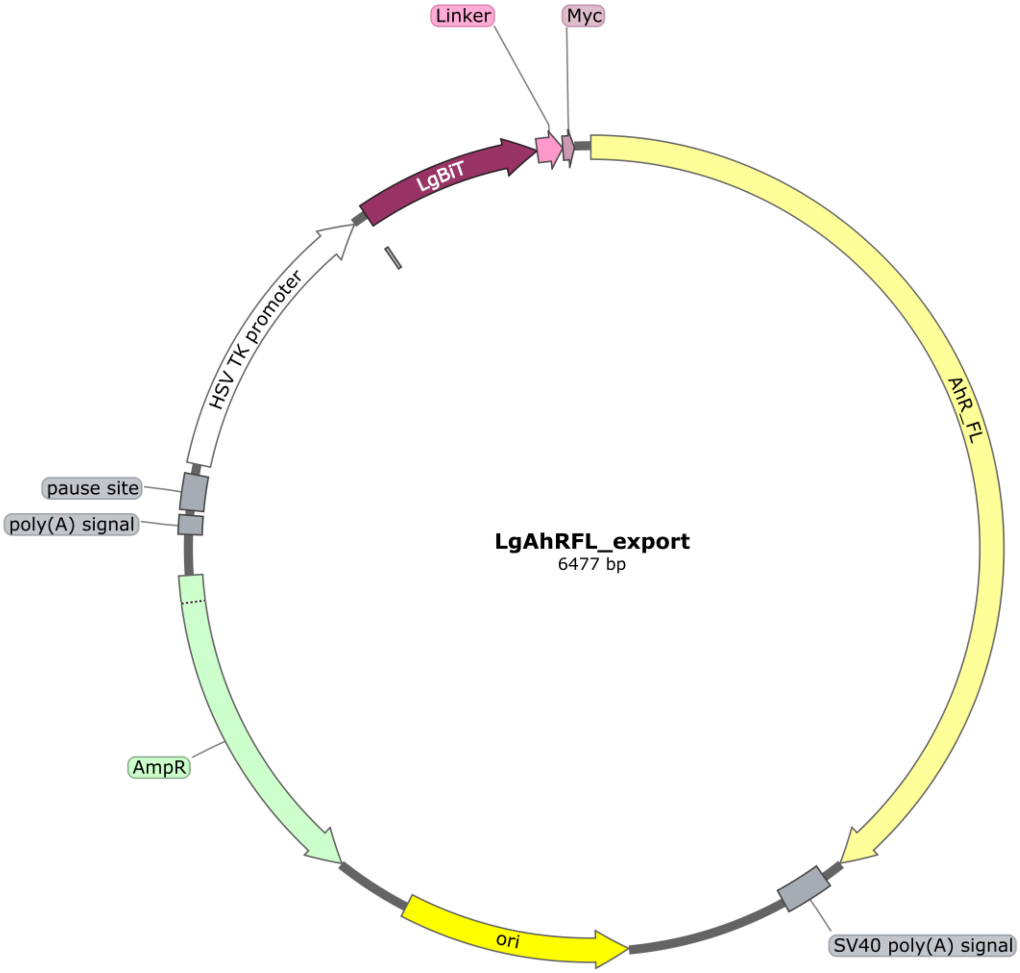

b) Vector map of NanoBiT ARNT plasmid.

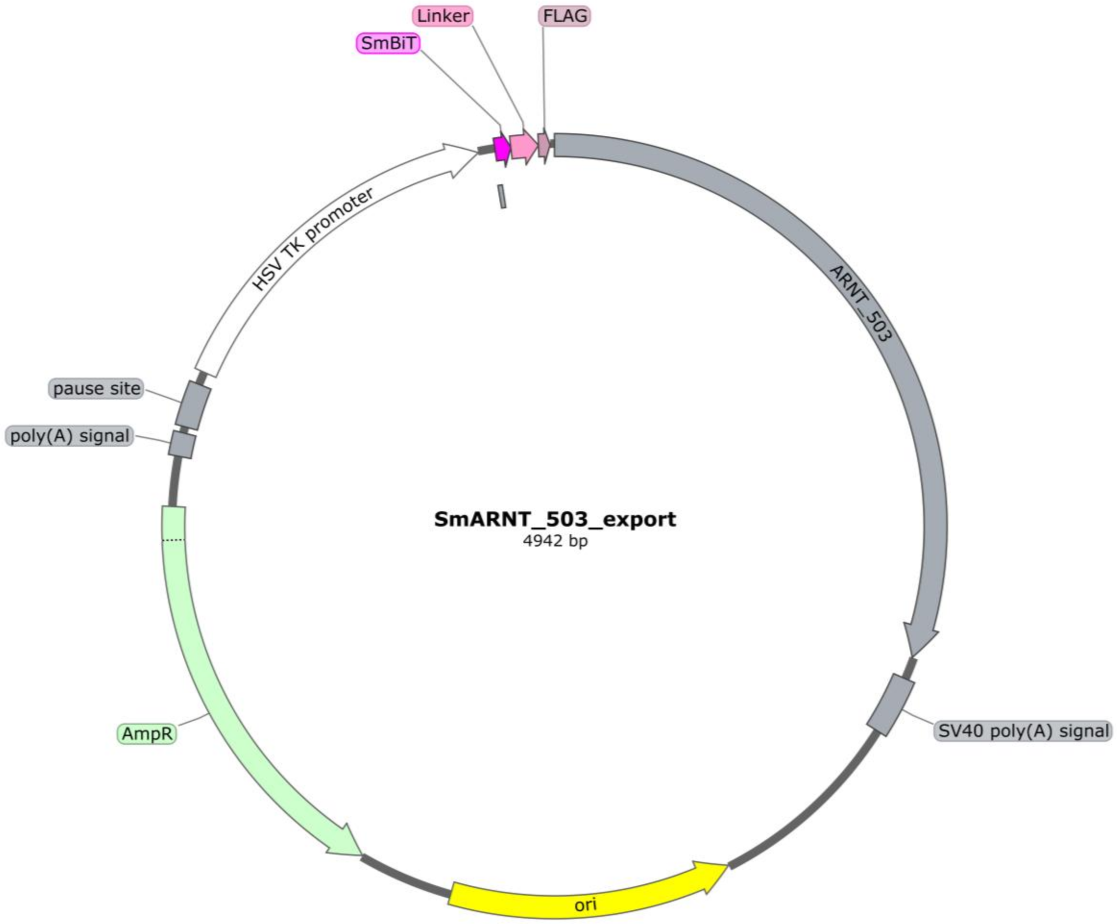

c) Vector map of NanoBiT HSP90 plasmid.

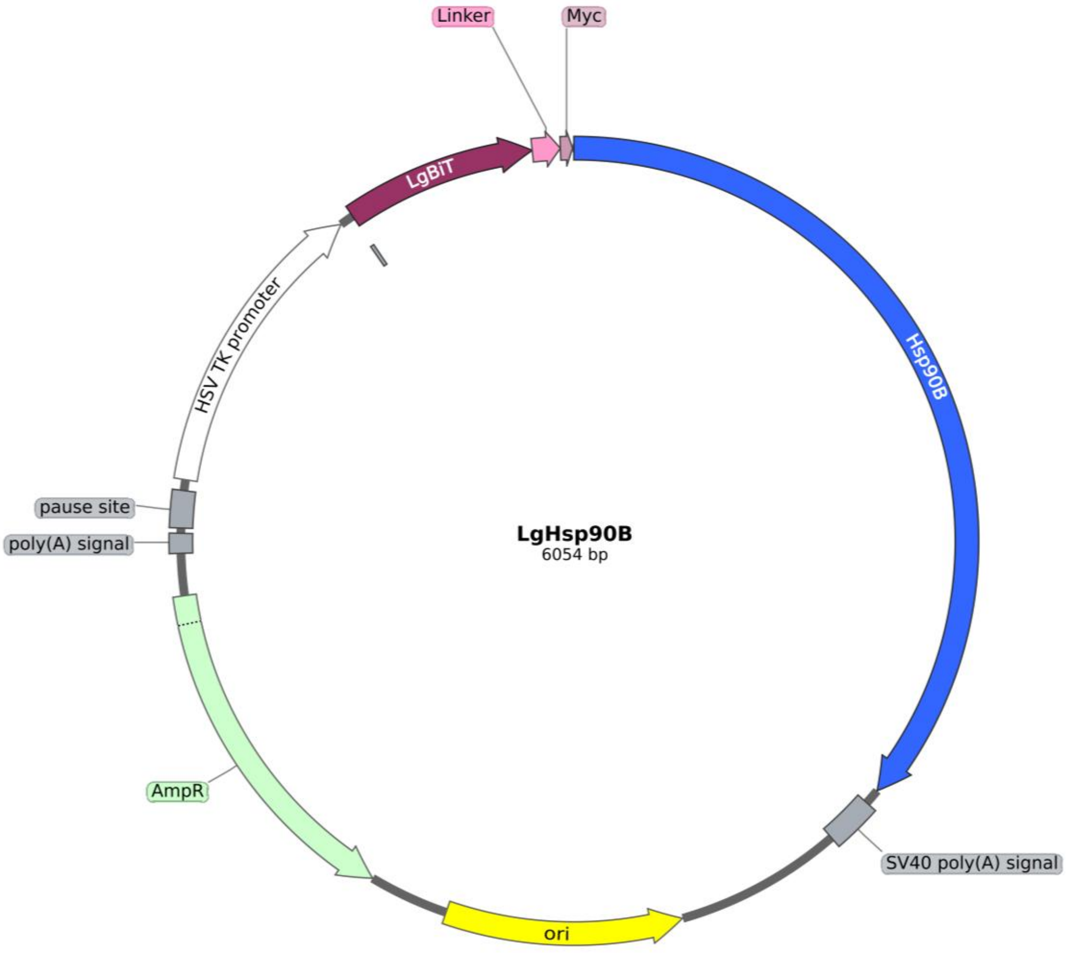

d) Vector map of BiFC AHR plasmid.

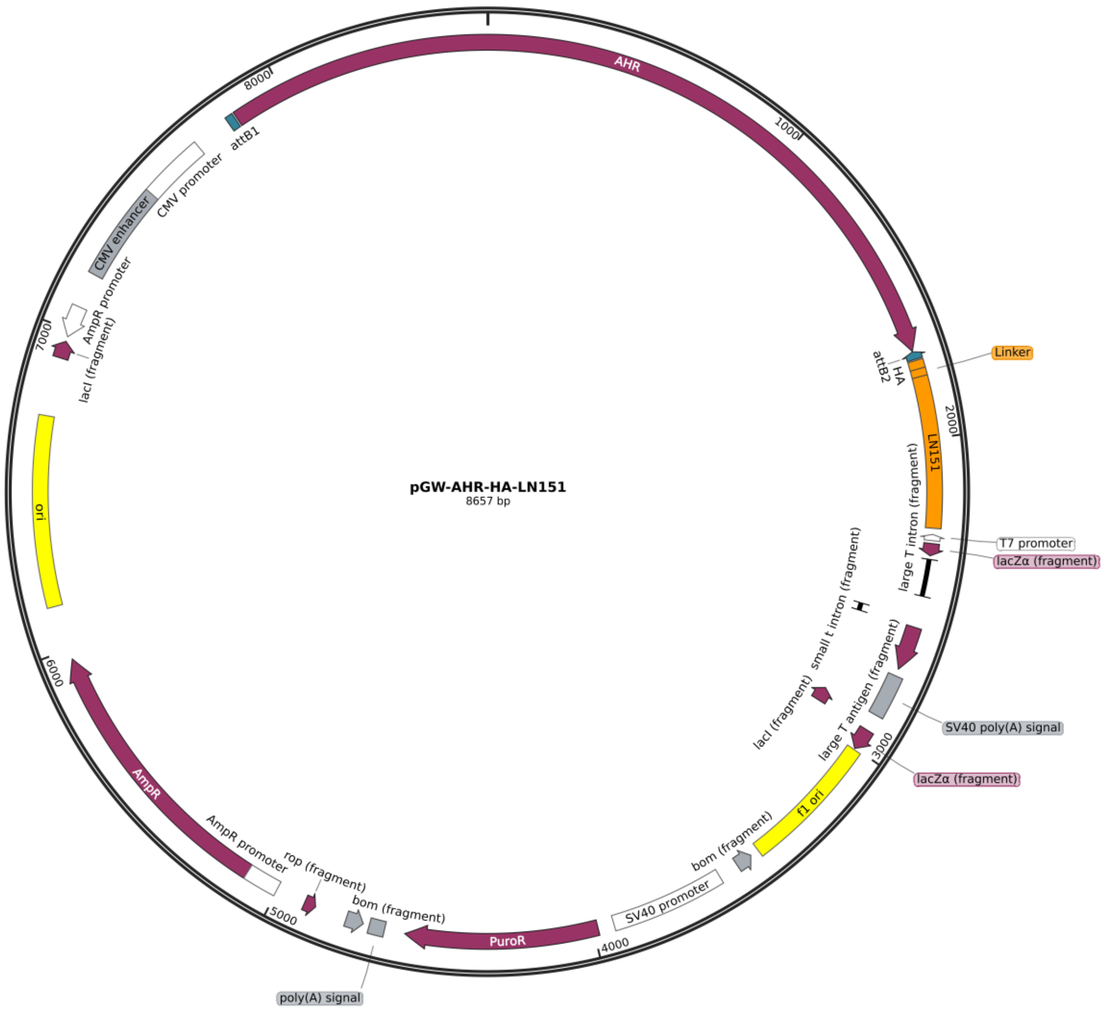

e) Vector map of BiFC ARNT plasmid.

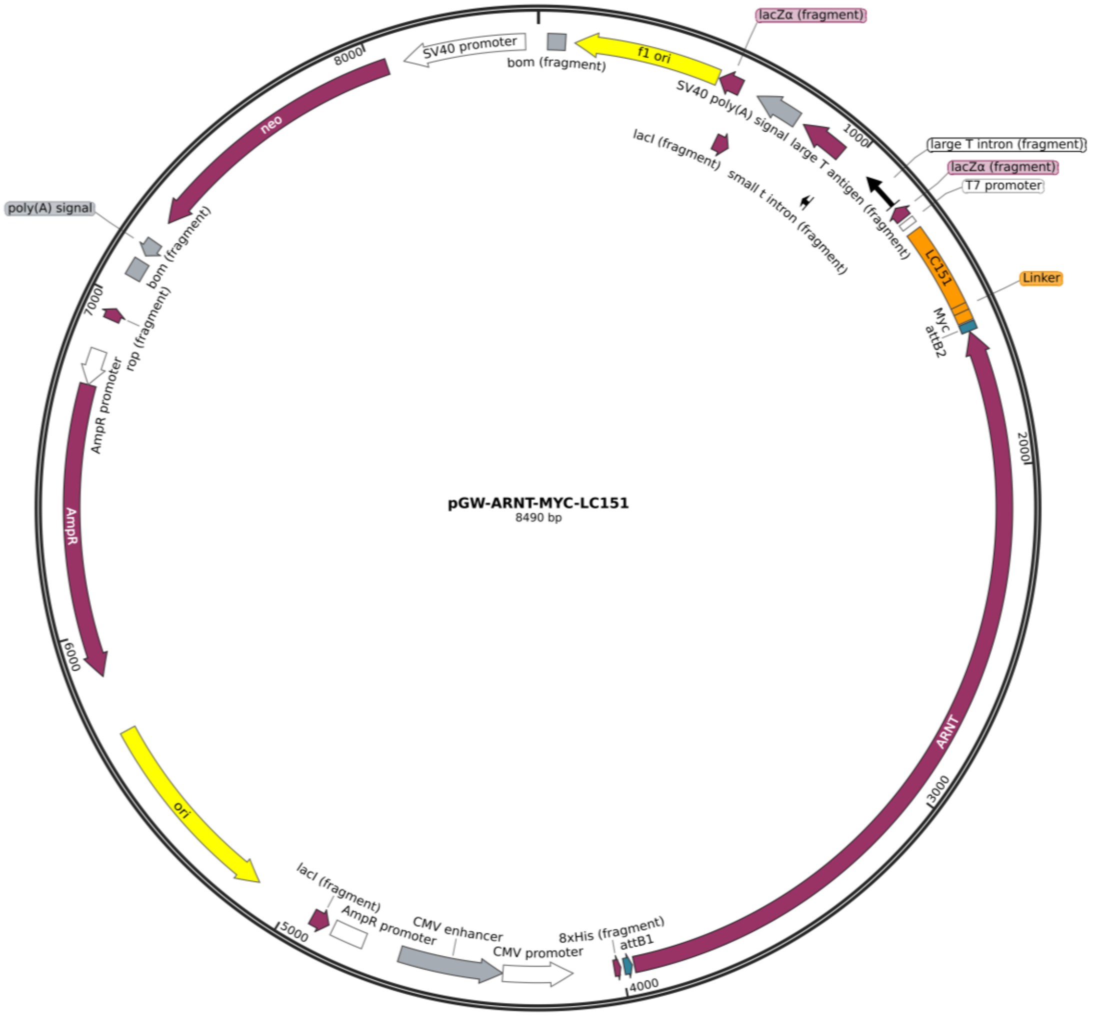

f) Vector map of BiFC HSP90 plasmid.

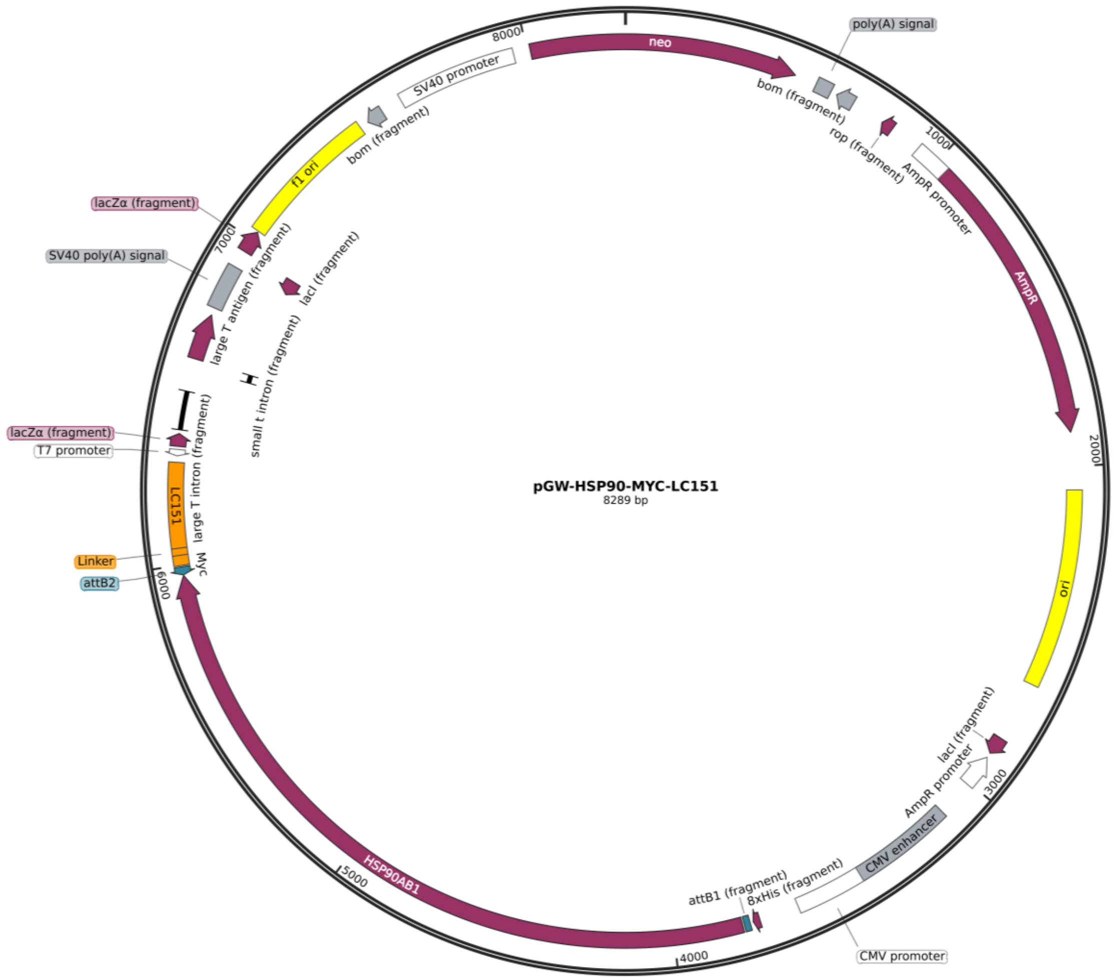
